# Tensile Expansion Mass Spectrometry for single cell metabolomics imaging

**DOI:** 10.64898/2026.08.15.745024

**Authors:** Jason A. Guerrero, Ethan A. Older, Marouen Zammali, Vignesh Venkataramani, Ramita Arampongpun, Danielle Latham, Dalia Riad, Jan Schwenzfeier, Alexander Potthoff, Diandra M. Vaval Taylor, Joanna E. Burdette, Roberto C. Andresen Eguiluz, Jens Soltwisch, Lydia Kisley, Laura M Sanchez

## Abstract

Matrix-assisted laser desorption/ionization mass spectrometry imaging (MALDI-MSI) enables the spatial mapping of endogenous biomolecules within native biological specimens; however, it remains limited in achieving single-cell resolution. While advances in instrument modifications, computational processing methods, and tissue-based sample preparation have facilitated high lateral resolutions and cellular level imaging, resolving metabolic heterogeneity at the single-cell level remains challenging for users without specific expertise or custom instrumentation. Here, we present tensile expansion mass spectrometry (TExMS), a cost-effective approach for single-cell MALDI-MSI that is compatible with commercial MSI instrumentation. TExMS utilizes highly stretchable hydrogels as a substrate for live-cell seeding, attachment, and desiccation, avoiding the need for chemical fixation and enabling the retention of both intracellular and extracellular metabolites, including media-derived components that are lost during fixation and washing. We used TExMS to expand individual cells of a human high-grade serous ovarian cancer (HGSOC) cell line and spatially map their small molecule (<800 Da) production. TExMS enabled ∼4-fold linear expansion of the hydrogel, translating to a ∼1.7-fold increase in average cell area and ∼1.3-fold increase in nuclear area and resulting in improved lateral resolution of metabolite distributions. Benchmarking against other platforms for high resolution MALDI-MSI, TExMS offered comparable spatial resolution to microgrid-enabled MALDI-MSI with 15 to 20-fold shorter acquisition times. We then used TExMS to map numerous intermediates from glycolysis, the tricarboxylic acid (TCA) cycle, and amino acid biosynthesis and probe the effects of serum starvation conditions on metabolic flux through these pathways, demonstrating a powerful use case for single-cell MALDI-MSI through TExMS.

**Table of Contents (TOC):** 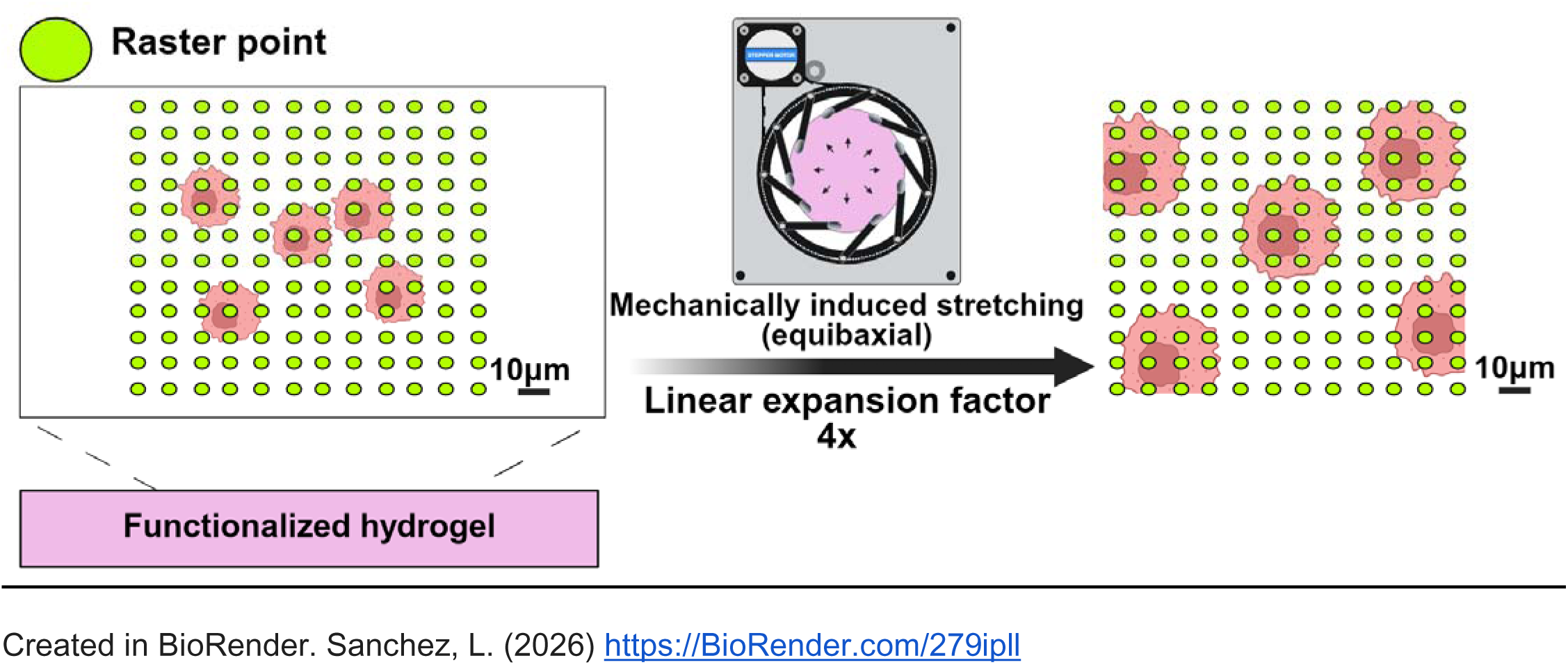

## Introduction

Small molecules play essential roles in biological systems, serving as metabolic building blocks, signaling molecules in cellular pathways, and regulators of diverse cellular processes. Determining the spatial location of these metabolites within subcellular compartments could provide important insights into the functional or dysfunctional states associated with diseases, such as in the cancer microenvironment. Cells interact with neighboring cells in a variety of manners including chemical signaling and nutrient exchange.

Even within the same tissue environment, individual cells are rarely biologically equivalent.^1^ Variability can arise from factors such as cellular age and microenvironmental differences.^1^ We will focus on discussing matrix-assisted laser desorption/ionization mass spectrometry imaging (MALDI-MSI)^2^ a technology, that allows for soft ionization, is a label-free analytical technique which enables high-throughput, sensitive detection of small molecules while providing high spatial information on the localization of analytes within broad range of biological specimens.^3,4^

MALDI-MSI has seen a number of changes due to its speed and high lateral spatial resolution (5-10 µm) capabilities. Since its inception, MALDI-MSI has undergone continuous technological advancements that have expanded the accessible mass range while improving resolving power and sensitivity.^5^ These developments have also driven efforts toward higher spatial resolution to achieve single-cell and even subcellular imaging. However, despite these advances, MSI remains both instrumentally and physically limited when applied to high-resolution single-cell imaging. One of the primary challenges in MALDI-MSI is its strong dependence on sample preparation. High-lateral resolution imaging requires careful optimization of preparation protocols to ensure reproducibility and sufficient ionization efficiency for reliable spectral acquisition.^6,7^ Commercial MALDI instruments are typically limited to a pixel size of approximately 5–10 μm, primarily due to constraints imposed by laser optics and the depth of focus. This presents challenges for single-cell analysis, as mammalian cells typically range from ∼10–100 μm in diameter. Further, smaller desorption plumes have less time for ion–molecule charge transfer processes because of limited radial expansion, ultimately leading to lower ion yields.^8,9^ There have been several approaches taken to realize single (sub)cellular metabolomics including hardware, computational, and sample preparation approaches.

Recent hardware modifications towards achieving single-cell MSI have largely focused on modifications to instrument design aimed to both increase lateral resolution while also keeping sensitivity in mind. As the area of measurement approaches subcellular levels; there are simply fewer moles of each analyte available for detection. One approach involves enhancing sensitivity through the addition of an orthogonal post-ionization laser, which interacts with the desorption plume and triggers a secondary ionization process. In this process, neutral analytes undergo charge transfer reactions with post-ionized matrix species, increasing sensitivity by up to two orders of magnitude.^10^ Another approach has been the development of transmission-mode MALDI (t-MALDI), which repositions the MALDI laser to irradiate samples from the backside while incorporating improvements in optical design, enabling ultra-high spatial resolutions with a pixel size approaching 1 μm or smaller (Fig. S1).^11,12^ However, the adoption of these advanced methods is often limited by accessibility. State-of-the-art MALDI instrumentation can be prohibitively expensive, and modifying existing instruments may involve significant costs and expertise.

Towards a computational level solution, software like SpaceM ^13^ or Fluorescence Integrated Single-Cell Analysis Script (FISCAS) ^14^ enables metabolic profiling across cells grown in 2D on glass slides to characterize cellular metabolic composition. FISCAS streamlines single-cell MALDI-MSI acquisition by automatically identifying cell-rich regions to minimize off-target during high spatial resolution MALDI-MSI experiments achievable with the state-of-the-instrumentation. In contrast, SpaceM computationally assigns ion signals to individual cells based on pixel overlap between the optical and ion images. As a result, metabolites signals are attributed to the entire cell, limiting its ability to resolve subcellular or organelle-level metabolic heterogeneity.^13,15^

Rather than modifying instrumentation, some groups have altered the samples themselves by building off the concepts introduced for expansion microscopy (ExM). ExM improves effective resolution through physical sample expansion using swellable hydrogels and has democratized microscopy workflows.^16^ In ExM, tissues or cells of interest are first chemically fixed and then permeabilized. The samples are then incorporated into an electronegative hydrogel precursor which is polymerized in and around the biological sample. The polymeric hydrogel matrix can be osmotically swelled to enlarge the sample to 4-20x its original size using common ExM protocols.^17–20^ Key to this process is the anchoring step that introduces reagents that bind target biomolecules, allowing them to remain tethered to the polymer network upon swelling. Structural components, such as the cell wall, are often partially or fully denatured during a digestion step to minimize mechanical constraints and prevent anisotropic distortion during osmotic expansion in water or low-salt buffers.^21^ Gel-assisted mass spectrometry imaging (GAMSI),^22^ tissue-expansion mass spectrometry imaging (TEMI),^23^ expansion imaging mass spectrometry (ExIMS)^24^ and tenfold expansion mass spectrometry imaging (10X ExMSI),^25^ are examples of ExM style sample based approach to achieve higher spatial resolution in MSI. However, these techniques differ in their analytical focus. GAMSI, ExIMS, 10X ExMSI have primarily been used to investigate the spatial distribution of native lipids in brain tissues—including phospholipids, sphingolipids, neutral lipids, and sterols—with proteins used mainly as a proof-of-concept demonstration.^22,24,25^ In contrast, TEMI aims to analyze a broader range of biomolecular classes in a variety of tissue types, including small molecules, lipids, peptides, proteins, and N-glycans.^23^ Although these methods are effective, they have been applied exclusively to tissues and only demonstrated with large Purkinje cells,^23^ placing limited emphasis on small molecules, individual cells, or subcellular organization.

Despite these advances, resolving metabolite distributions at the single cell and subcellular level remains challenging. To address this challenge, we have developed tensile expansion mass spectrometry (TExMS), an inexpensive method that is compatible with commercial MALDI instrumentation. Using TExMS, live cells in 2D culture are seeded on top of a functionalized hydrogel substrate and allowed to adhere.

Following adherence, the cells are subjected to controlled multiaxial mechanical expansion through the application of tensile force using an iris expansion device resulting in the physical expansion of live cells. Since TExMS does not require fixation, gelation, or osmotic swelling, our data highlight that it promotes the retention of metabolites that are often lost during conventional sample preparation workflows, although it requires a more extensive sample preparation process. With this tensile expansion platform, we effectively increase the number of spatial sampling points per cell during MSI acquisition, enabling higher-resolution metabolomic mapping of individual cells. We compared the TExMS method to t-MALDI using the OVCAR8 high-grade serous ovarian cancer cell line, to map the different metabolites in the glycolysis and TCA pathways and compare the results to serum starvation to validate that metabolic changes are visible at the subcellular level.^26^

## Results & Discussion

To begin the development of TExMS, we focused on the first principles required to achieve ionization and reproducible MSI data acquisition, including functionalization of the chemically inert hydrogel surface, establishment of 2D cell adhesion, visualization of OVCAR8 cellular response to mechanical stretching, and preparation of a MALDI-compatible sample for analysis (Figure 1). A critical component in tensile expansion is allowing the living cells to adhere themselves to the hydrogel surface to withstand the physical force of the stretching. We purposely avoided degradation of the cell wall and chemical fixation in our development of TExMS to preserve the distribution of metabolites and lipids. A general workflow for TExMS is shown in Figure 1.

**Figure 1:**
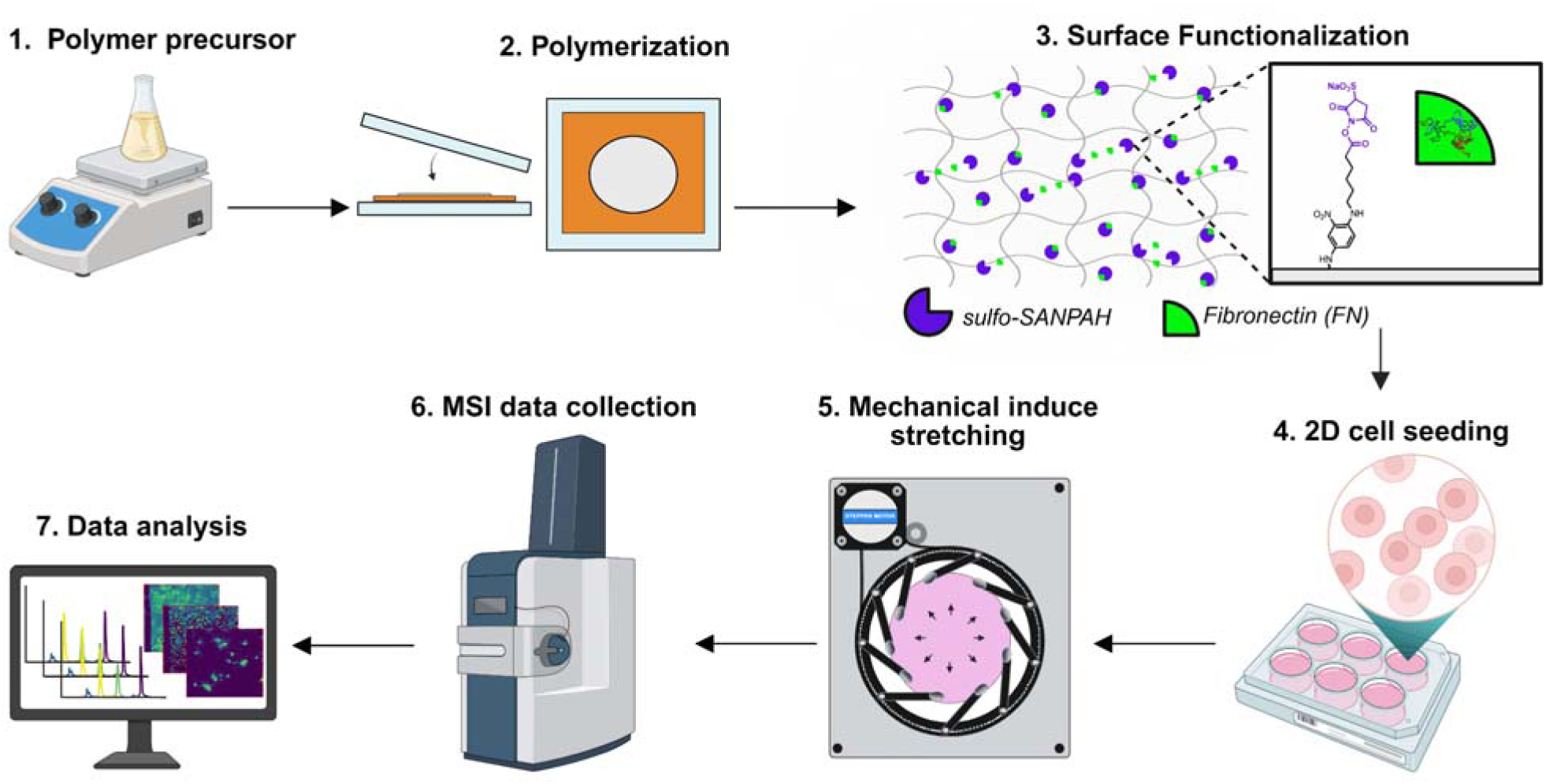
Generalized workflow of TExMS. **(1)** Hydrogel monomer stock solutions were prepared at concentrations of 2.91% (w/w) alginate (Alg), 28.57% (w/w) acrylamide (AAm), 1.96% (w/w) N,N′-methylenebisacrylamide (MBA), and 9.1% (w/w) ammonium persulfate (APS). The hydrogel precursor solution was prepared by combining the stock solutions to final concentrations of 1.8% (w/w) Alg, 10.62% (w/w) AAm, with MBA and APS at 0.073% and 0.58% (w/w), respectively, relative to the weight of AAm. The resulting solution was stirred for 45 minutes. N,N,N′,N′-Tetramethylethylenediamine (TEMED) was then added to a final concentration of 0.61% (w/w) and mixed for an additional 15 minutes before casting. The final hydrogel contained a total polymer concentration of 12.42% (w/w) (Alg + AAm). **(2)** Following homogenization with TEMED, glass slabs were primed with methanol (MeOH) to secure polyethylene terephthalate (PET) sheets. A silicone mold was placed between the glass slabs and carefully pressed to eliminate trapped air that could interfere with polymerization. The hydrogel precursor solution was poured into the mold, sealed with a second PET-lined glass slab, and incubated at 35°C for 5 hours, followed by overnight equilibration at room temperature. The hydrogels were then removed from the mold and immersed in a 20 mM CaSO_4_·2H_2_O slurry to ionically crosslink the Alg network, producing a highly stretchable alginate-Ca²□/polyacrylamide DN hydrogel. **(3)** The hydrogel surface was functionalized with sulfo-SANPAH, a heterobifunctional photo-crosslinker containing a photoactivatable nitrophenyl azide group and an amine-reactive N-hydroxysuccinimide (NHS) ester. This chemistry enabled covalent immobilization of fibronectin (FN), generating an artificial ECM that promotes cell adhesion. **(4)** Following functionalization with fibronectin, OVCAR8-RFP cells were seeded onto the hydrogel surface and cultured for 24 hours at 37°C in a humidified atmosphere containing 5% CO_2_ to allow cell attachment and spreading. **(5)** Hydrogels exhibiting appropriate cell morphology and adhesion were subjected to controlled multiaxial mechanical stretching to physically expand the sample. The expanded hydrogels were subsequently desiccated for 4–6 hours and coated with MALDI matrix in preparation for MSI analysis. **(6,7)** MALDI-MSI data was acquired using a Bruker timsTOF flex mass spectrometer, followed by downstream visualization, processing, and analysis using SCiLS Lab software. Created in BioRender. Sanchez, L. (2026) https://BioRender.com/0l5pbcm

### Functionalization of hydrogel surface for 2D cell adhesion

A double network (DN) hydrogel is used as an expandable substrate because its combination of covalent and ionic crosslinking enables large deformation without fracture. In this biocompatible system, Ca^2+^- crosslinked alginate acts as a sacrificial ionic network that dissipates energy under strain, while covalently crosslinked polyacrylamide maintains elasticity and structural integrity of the whole hydrogel. Together, these networks provide the hydrogel with high stretchability and toughness.^27^ The DN hydrogel is chemically inert, some cell lines are able to undergo 2D cell adhesion without surface functionalization.^28^ Initially, the DN hydrogel did not support proper 2D cell adhesion or normal cellular morphology in the absence of surface functionalization. Without functionalization, murine ovarian surface epithelial (MOSE), murine oviductal epithelial (MOE), and human OVCAR8-RFP epithelial cells were unable to properly and effectively adhere to the hydrogel surface. Therefore, optimization of the hydrogel surface was performed to enable effective 2D cell adhesion of the OVCAR8-RFP (red fluorescent protein) epithelial cells.^29^ The OVCAR8-RFP cell line was selected due to its rapid *in vivo* peritoneal metastasis as an ovarian cancer model and stable expression of cytosolic red fluorescent protein.^30^ To improve cell adhesion to the surface functionalization, a sulfo-SANPAH protocol was adapted and optimized.^31^ Sulfo-SANPAH acts as a heterobifunctional crosslinker that enables covalent attachment of extracellular matrix (ECM) proteins to otherwise chemically inert surfaces, thereby facilitating stable 2D cell adhesion through the introduction of anchored surface adhesion proteins. Due to their relatively large size, the hydrogels were trimmed to 30 mm, referred to as hydrogel plugs herein, to fit in 6 well plates (30 mm diameter) required for subsequent cell culture. The protocol described by Smith *et al*. was modified to our sample sizes prior to UV photolysis to ensure complete surface coverage.^31^ This optimized functionalization enabled OVCAR8-RFP cells to adopt a polygonal morphology characteristic of adherent mammalian epithelial cells cultured under standard tissue culture conditions (Fig. S2). Cell adhesion to the hydrogel substrate was markedly improved, with some cells exhibiting partial embedding within the hydrogel network. Importantly, the cells remained attached to the substrate following mechanical stress testing. Overnight incubation further demonstrated continued cell proliferation, confirming that the functionalized hydrogel surface supported both stable cell adhesion and sustained growth (Fig. S3).

Fibronectin (FN) was selected as the ECM protein for hydrogel functionalization because it is a major component of the extracellular matrix encountered by OVCAR8-RFP cells during metastasis. FN functionalization resulted in robust adhesion of OVCAR8-RFP cells to the hydrogel substrate. Several FN concentrations were evaluated and the optimal condition was determined to be 350 μg/mL in MilliQ H_2_O (Fig. S4). With FN, OVCAR8-RFP cells adhered to the hydrogel and formed stable attachment to the surface.

During the initial implementation of the sulfo-SANPAH functionalization different incubation temperatures ranging from 4 °C to 37 °C were also evaluated. A noticeable improvement in cellular morphology and adhesion was observed when FN-coated hydrogels were incubated at 4 °C for 24 hours (Fig. S5). The colder hydrogel incubation temperature resulted in polygonal cellular morphology and more consistent adhesion across the entire hydrogel surface. To further ensure that OVCAR8-RFP cells adhered specifically to the central region of the functionalized hydrogel substrate, a dry-seeding approach was employed.

### Visualization of OVCAR8-RFP under mechanical expansion

Prior to mechanical stretching, hydrogels were mounted onto a custom 3D-printed expansion apparatus (iris) consisting of two components that were assembled to secure the hydrogel at nine equally spaced stress points, ensuring uniform multiaxial stretching (additional details are provided in the Methods section).^28^ After the tensile expansion, hydrogels containing OVCAR8-RFP cells were secured using drying rings to maintain their expanded state prior to desiccation and MALDI sample preparation. Under both brightfield and fluorescence imaging, OVCAR8-RFP cells were readily observed prior to tensile expansion. Following expansion, it was more challenging to clearly distinguish individual cells on the hydrogel surface in brightfield imaging; however, fluorescence imaging allowed visualization of cytosolic-RFP cells adhered to the hydrogel substrate. Although the bulky nature of the iris drying rings used to secure the gel after stretching limited our ability to obtain high resolution images after expansion, the detection of a robust cytosolic-RFP signal suggested that the cell membrane remained intact after expansion. If the cells had ruptured, we would have expected that the RFP would leak from the cytosol and be diluted into the background hindering detection.

The theoretical maximum expansion derived from the geometry of the device, that the current version of the iris expansion device can apply onto the hydrogel, is ∼3.8x. The fact that the hydrogels need to be large enough to load onto the device and the device needs to be small enough to fit onto a microscope limits the expansion factor. Engineering tolerances and softness of the hydrogels also have an impact on the maximum expansion factor realized by a sample. To orthogonally validate that tensile stretching increased the spatial dimensions of cells, the area of OVCAR8-RFP cells was manually measured pre- and post- expansion.

Hydrogels containing two-dimensionally adhered OVCAR8-RFP cells were treated and stained with the Hoechst nuclear stain to enable visualization of nuclei (Fig. S6). Under these conditions, both cellular and nuclear areas were quantified. Pre-expanded cells exhibited relatively uniform sizes with low cell-to-cell heterogeneity and an average cellular area of approximately 349 ± 110 µm^2^ (Figure 2a,b) Following tensile expansion, the cells retained their overall morphology and appeared noticeably larger with a statistically significantly larger average area of approximately 1046 ± 366 µm^2^ (Figure 2a,b). Using the Hoechst stain, we used a particle analysis to automatically measure the average nuclear area of the cells, finding that it significantly increased from 148 ± 72 µm^2^ in pre-expanded cells to 255 ± 109 µm^2^ in post-expanded cells (Figure 2a,c). The 3.0-fold increase in the average cellular area translates to a ∼1.7x linear expansion factor and the 1.7- fold increase in average nuclear area translates to a ∼1.3x linear expansion factor. These results suggest that while the cells are indeed larger after expansion, their adhesion to the hydrogel surface may be limiting their potential expansion as evidenced by the total linear expansion factor observed compared to the theoretical maximum linear expansion factor of 3.8x. Further, the difference in linear expansion factor between cellular area and nuclear area may indicate a differential application of tensile expansion to the overall cell versus the nucleus. However, this difference may also be derived from the manual measurement of the cellular area compared to the particle analysis measurements of their nuclear area. The observed linear expansion of the nucleus also suggests that tensile expansion is applied not only to the cellular membrane but throughout the cell, without the use of chemical anchoring, showing the potential and promise of TExMS.

**Figure 2:**
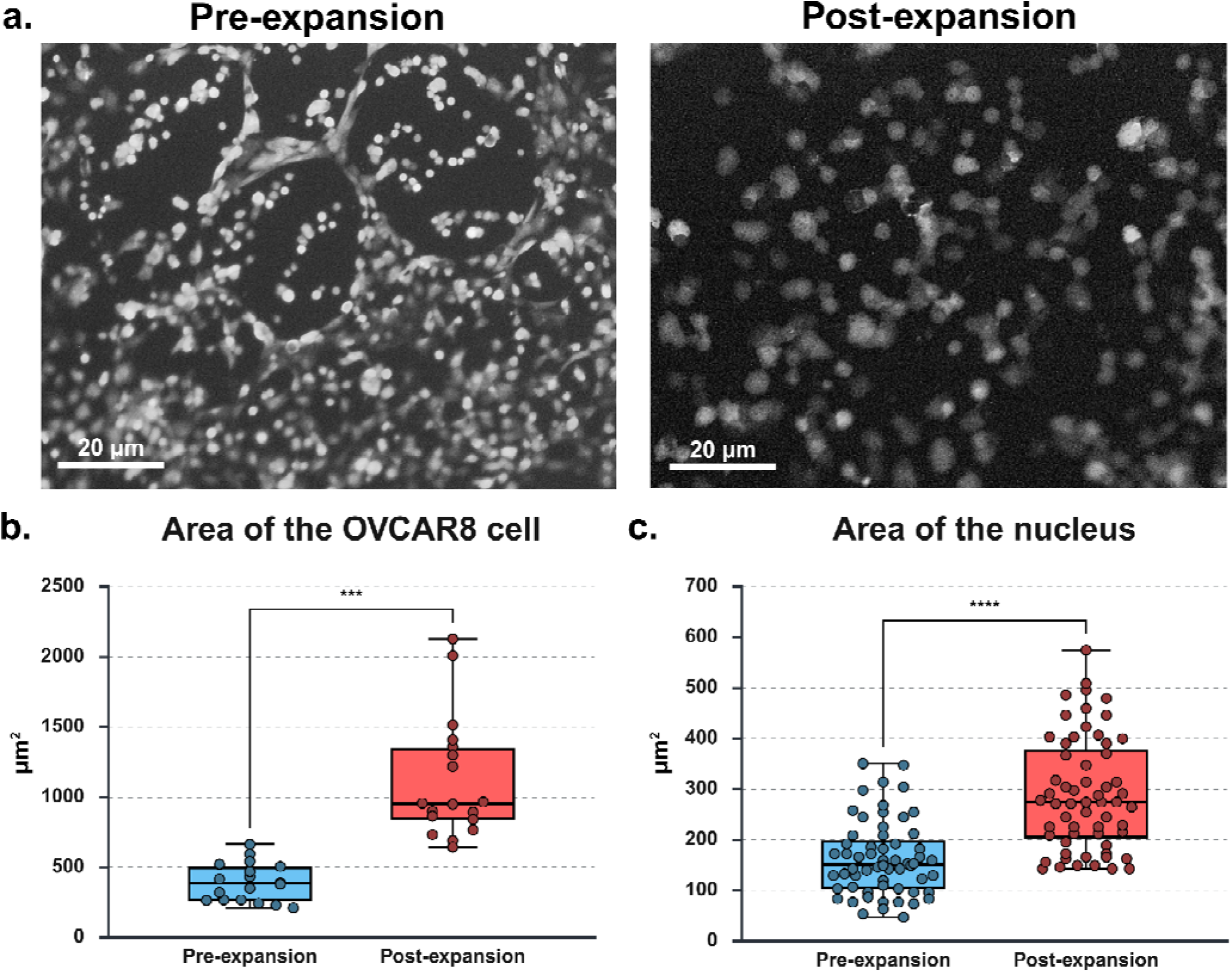
Evaluation of OVCAR8-RFP cell and nuclear area pre- and post-expansion. **(a)** Representative fluorescence images of pre- and post-expanded OVCAR8-RFP cells and outlined. Hydrogels containing adhered cells were subjected to tensile multiaxial stretching, after which cell and nuclear areas were quantified using ImageJ. Measurements were obtained from four independently prepared hydrogels. An unbiased subset of cells was selected for analysis. Statistical comparisons between pre- and post-expanded conditions were performed using a paired t-test and a nonparametric test. A statistically significant increase in **(b)** cellular area was observed following expansion (p-value of 1.02 × 10^-5^). Similarly, a significant increase in **(c)** nuclear area was detected (p-value of 1.06 × 10^-11^), confirming that mechanically induced stretching effectively increases both whole-cell and nuclear dimensions. Created in BioRender. Sanchez, L. (2026) https://BioRender.com/nivrcmt

### Optimization of TExMS for MSI compatibility

MALDI can be prone to ion suppression from the biological matrices and consistent sample height ensures reproducibility of signals within and across experiments. We have pursued many different sample preparation techniques and protocols and provide a detailed explanation of the results in the Supporting Information. To ensure the hydrogel remained within the acceptable height tolerance of the timsTOF fleX (±70 μm) beyond which laser focus, mass accuracy, and ionization efficiency can be affected, we applied 18 mm diameter coverslips with thicknesses ranging between 0.13-0.17 mm to underside of the hydrogels during desiccation. The hydrogels were held in place using magnetic drying rings and dried for 4–6 hours at room temperature. Once fully dried, the samples were trimmed and the exposed coverslip and sample edges were permanently secured onto glass slides using a clear cyanoacrylate adhesive, producing a flat and stable surface suitable for MALDI-MSI acquisition.

### Comparison of TExMS with other single cell methods

To compare the ionization efficiency of TExMS with existing MALDI-MSI methodologies, four different analyses were performed: 1) cell fixation on poly-D-lysine (PDL) coated glass slides followed by analysis on a timsTOF fleX (condition 1), 2) analysis using a timsTOF fleX with microgrid capabilities (condition 2), 3) analysis on a state of the art, modified transmission mode timsTOF fleX (condition 3), and 4) unfixed cells adhered to hydrogels via TExMS and analysis on timsTOF fleX. All samples were subjected to matrix application either by sublimation on a Bruker superlimator or by spraying using an HTX TM-Sprayer prior to analysis (Figure 3a–d). N-(1-naphthyl)ethylenediamine dihydrochloride (NEDC) was the matrix of choice due to its compatibility with dual polarity detection and small-molecule focus.^32^ TExMS-specific samples (Figure 3d) were prepared under conditions comparable to the 10 µm pixel size workflow (condition 1 described above, Figure 3a), with the exception of increased laser shots (1000 shots) per pixel. At the theoretical maximum expansion factor of the iris device, we expect ∼3.8-fold expansion which would translate to an effective pixel size of ∼2.6 µm^2^ after expansion. When measuring the average cellular and nuclear areas, we observed linear expansion factors of ∼1.7x and ∼1.3x respectively. Based on the average cellular area, this suggests that the effective pixel size we realized was ∼5.9 µm^2^. However, there may be other factors influencing the overall linear expansion factor realized by the cells and a biologically independent marker would be required to more accurately determine the actual linear expansion factor affected upon the cells. To improve surface uniformity during desiccation and maintain optimal focal alignment at high spatial resolution, a circular coverslip is placed beneath the sample to reduce warping and improve reproducibility.

**Figure 3:**
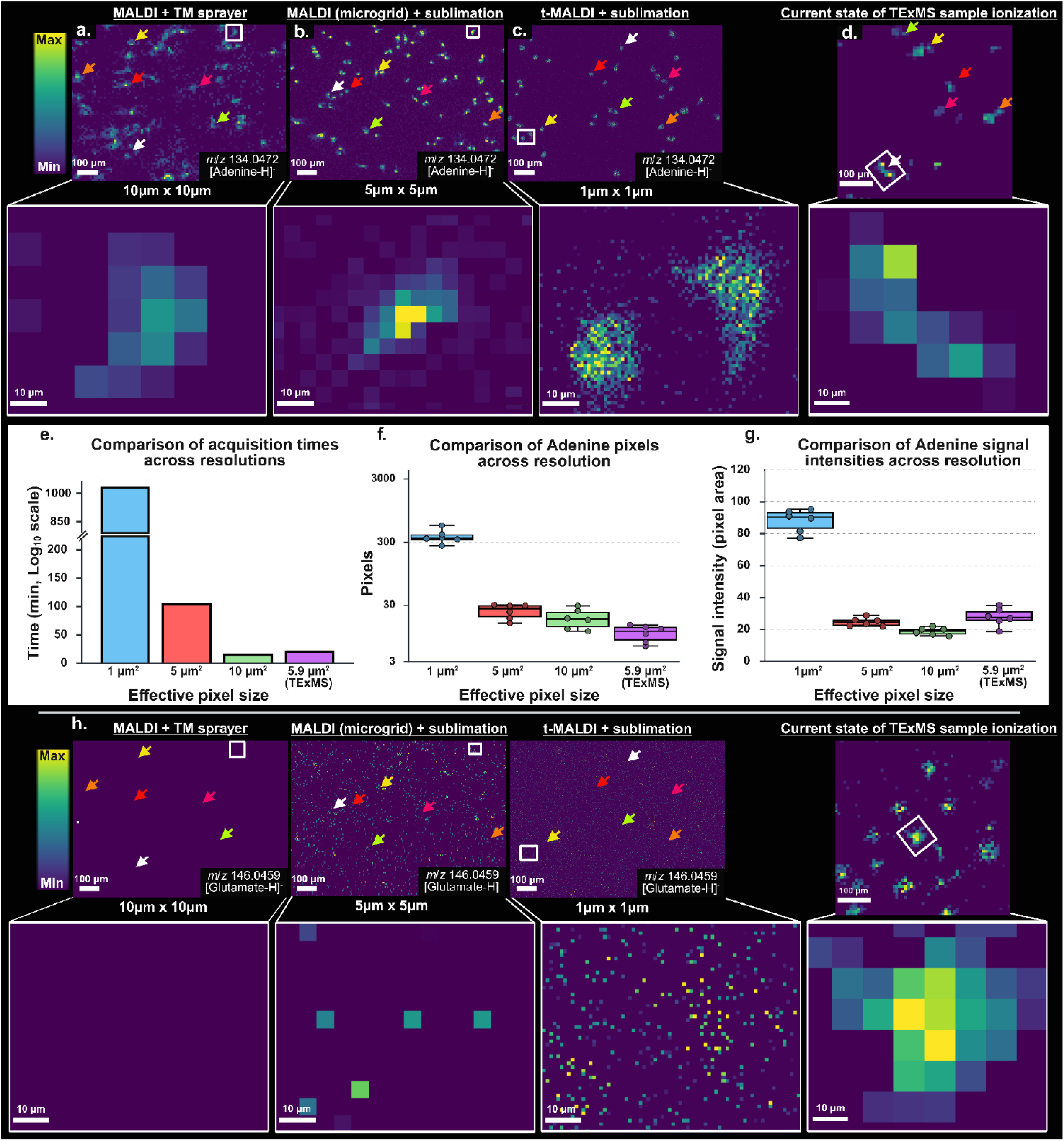
The current state of single-cell MALDI-MSI and development of TExMS. All data presented in this figure were acquired using a Bruker timsTOF fleX instruments that provides a resolving power of ∼65,000 FWHM at *m/z* 1222 in both positive and negative ion modes. **(a–c)** Current state of single-cell imaging using fixed OVCAR8-RFP cells deposited on glass slides, tracking the highly ionizable ion at *m/z* 134.0472 ([adenine–H]^−^) in negative ion MS mode. **(a)** Samples were coated with NEDC matrix (90:10 ACN:H_2_O) using an HTX TM-Sprayer. Data were acquired using a custom single-spot configuration (scan range: 6 × 6 μm; field size: 10 × 10 μm) with a laser power of 85%, 5,000 Hz frequency, single burst, and 500 shots. Total acquisition time was 1 hour and 9 minutes. A zoomed region (white) highlights a single cell; however, at 10 μm spatial resolution, subcellular localization remains difficult to resolve. **(b)** Imaging at 5 μm resolution was achieved using a Bruker microgrid. NEDC matrix was dissolved in 100% MeOH, sublimated, and recrystallized with 0.5% EtOH at 65 °C for 150 seconds. Laser parameters included a scan range of 6 × 6 μm, field size of 5 × 5 μm, 80% laser power, 10% global attenuator, 1,000 Hz frequency, single burst, and 25 shots. Total acquisition time was 6 hours 13 minutes. **(c)** Transmission-mode MALDI (t-MALDI) enabled ∼1 μm spatial resolution, providing enhanced localization of adenine compared to 5–10 μm datasets. Norharmane matrix (100% EtOH) was applied and recrystallized under the same conditions as in **(b)**. Laser settings included 80% power, 10,000 Hz frequency, single burst, and 15 shots, with a total acquisition time of 18 hours and 16 minutes. **(d)** TExMS sample matrix application was performed under conditions similar to those used for 10 µm spatial resolution, with the exception that the number of laser shots was increased to 1000 shots. The total data acquisition time for the run was approximately 45 minutes. Adenine, *m/z* 134.0472 ([M–H]^−^) was used as a reference marker during MALDI sample preparation to evaluate ionization efficiency and spatial consistency. **(e)** Comparison of average time for each data acquisition (graph displayed on a log_10_ scale). **(f-g)** Six individual cells were selected and annotated with colored arrows—cell 1 (orange), cell 2 (red), cell 3 (yellow), cell 4 (white), cell 5 (magenta), and cell 6 (green)—to enable direct comparison across spatial resolutions and the TExMS method. **(f)** Pixel-level analysis demonstrates that higher spatial resolution preserves cellular morphology, allowing clear delineation of individual cell boundaries, whereas lower-resolution data results in a loss of morphological detail and spatial definition. **(g)** The average signal intensities across all conditions were quantified and compared to evaluate relative ionization efficiency. **(h)** The TExMS workflow avoids fixation and extensive rinsing steps that can contribute to small-molecule delocalization. Because small molecules are not covalently anchored within the sample matrix, they can diffuse out during conventional single-cell preparation, resulting in the loss of metabolites such as L-glutamate. In contrast, the L-glutamate signal is retained in TExMS, demonstrating the workflow’s ability to preserve small-molecule metabolites that may otherwise be lost during conventional sample preparation. Created in BioRender. Sanchez, L. (2026) https://BioRender.com/bm2ygq0

The total acquisition time for the TExMS run was approximately 45 minutes which was orders of magnitude less than the MALDI-microgrid (condition 2) and t-MALDI (condition 3) experiments. The highly ionizable ion at *m/z* 134.0472 corresponds to adenine was used ([M–H]) as a reference marker to assess ionization efficiency and spatial consistency across these four experiments to compare and contrast acquisition time and performance across spatial resolutions (Figure 3e-g). At 1 µm pixel size (t-MALDI, condition 3), subcellular structures were highly resolved, but acquisition time increased approximately 10–20-fold relative to the 5-10 pixel size workflows (conditions 1, 2). Although higher spatial resolution generated a greater number of pixels and improved morphological definition, it also resulted in reduced signal intensity (Figure f,g), likely due to the smaller laser spot size and reduced sampled volume per pixel, which decreases ion yield and overall ionization efficiency. To quantify these effects, six individual cells were analyzed per condition for direct comparison across spatial resolutions and the TExMS workflow. Pixel-level analysis demonstrated that higher spatial resolution preserves cellular morphology and enables clear delineation of cell boundaries, whereas lower-resolution imaging results in reduced spatial definition (Figure 3a–d). Average signal intensities were normalized to the effective pixel area, with TExMS data accounting for the smaller effective pixel size given by the observed linear expansion of ∼1.7x, across all experimental conditions to evaluate relative signal intensity and compare the performance of conventional MALDI-MSI with the TExMS workflow. Normalization by pixel area enabled direct comparison of signal intensities between methods despite differences in physical dimensions. Strikingly, when compared with 5 and 10 μm pixel sizes in conventional MALDI-MSI, TExMS exhibited higher signal intensity per unit area. This improvement is attributed to the tensile stretching process, which physically expands the sample, increasing the effective sampling area while taking advantage of the reduced acquisition time for 10 μm acquisitions.

This approach enables improved detection of metabolites such as L-glutamate, which are otherwise susceptible to diffusion and loss during conventional sample preparation (Figure 3h). Traditional single-cell MALDI-MSI workflows, including t-MALDI (Fig. S7), rely on cell fixation to achieve improved lateral spatial resolution. However, fixation and subsequent washing steps can lead to analyte delocalization and the loss of small, highly diffusible metabolites, resulting in the loss of biologically relevant molecular information prior to analysis. In contrast, TExMS utilizes live cells that are subsequently desiccated on a hydrogel substrate, promoting the retention of small molecules while preserving their spatial localization. Furthermore, despite requiring a more extensive sample preparation workflow, TExMS substantially reduces data acquisition time (Figure 3e) while remaining compatible with conventional MALDI imaging instrumentation. These results highlight TExMS as a promising advancement in expansion approaches by providing a balance between spatial resolution and ion signal preservation that is comparable to conventional MALDI imaging workflows.

Collectively, these findings support the use of TExMS to advance imaging strategies for single-cell met analysis. bolite

### TExMS maps metabolic responses to cellular stress and nutrient availability

Leveraging TExMS and the matrix NEDC which efficiently ionizes small molecules such as intracellular and media-derived compounds, we sought to map metabolic intermediates in OVCAR8-RFP cells. We first constructed an in-house metabolome library for the OVCAR8-RFP cell line using high resolution LC-MS/MS and database matching with GLobal Natural Products Molecular Networking 2 (GNPS2)^33^ and the Human metabolome Database (HMDB).^33,34^ We annotated metabolites associated with the TCA cycle, including amino acids, organic acids, glucose, and lipids (Table S1). This in-house library enabled us to map numerous metabolic intermediates in single OVCAR8-RFP cells using complementary target spatial metabolomics approaches focused on glycolysis and the TCA cycle (Figure 4). Using t-MALDI on unexpanded cells grown on glass slides, we could annotate a few metabolic intermediates and end products including ATP, and AMP (Figure 4a and S1). Using TExMS, we annotated key metabolic intermediates in glycolysis including glucose-6- phosphate and pyruvate, the first and final regulatory steps in the pathway, respectively (Figure 4b). In addition, we annotated glycerol 3-phosphate, an important precursor to glycerolipid biosynthesis, and several amino acids including branched-chain amino acids which we have previously shown to be dysregulated in ovarian cancer cells to fuel rapid growth (Figure 4b).^35^ Among TCA cycle intermediates, we could annotate citric acids, isocitric acid, α-ketoglutarate, succinic acid, and malic acid (Figure 4c). Interestingly, α- ketoglutarate was readily detected by TExMS, suggesting its increase abundance compared to other TCA cycle intermediates which is a potential manifestation of the Warburg effect, in which the TCA cycle is dysregulated in favor other metabolic pathways such as amino acid biosynthesis.^36,37^ We also observed clear co-localization of succinic acid and malic acid which was segregated away from a putative membrane-associated lipid, suggestive of the mitochondrial matrix containing TCA cycle intermediates away from the cell membrane and demonstrating that TExMS allows subcellular detection of important metabolic intermediates (Figure 4d). Compared to our analysis using t-MALDI, TExMS allowed us to improve our metabolite coverage over glycolysis, amino acid biosynthesis, and the TCA cycle.

**Figure 4:**
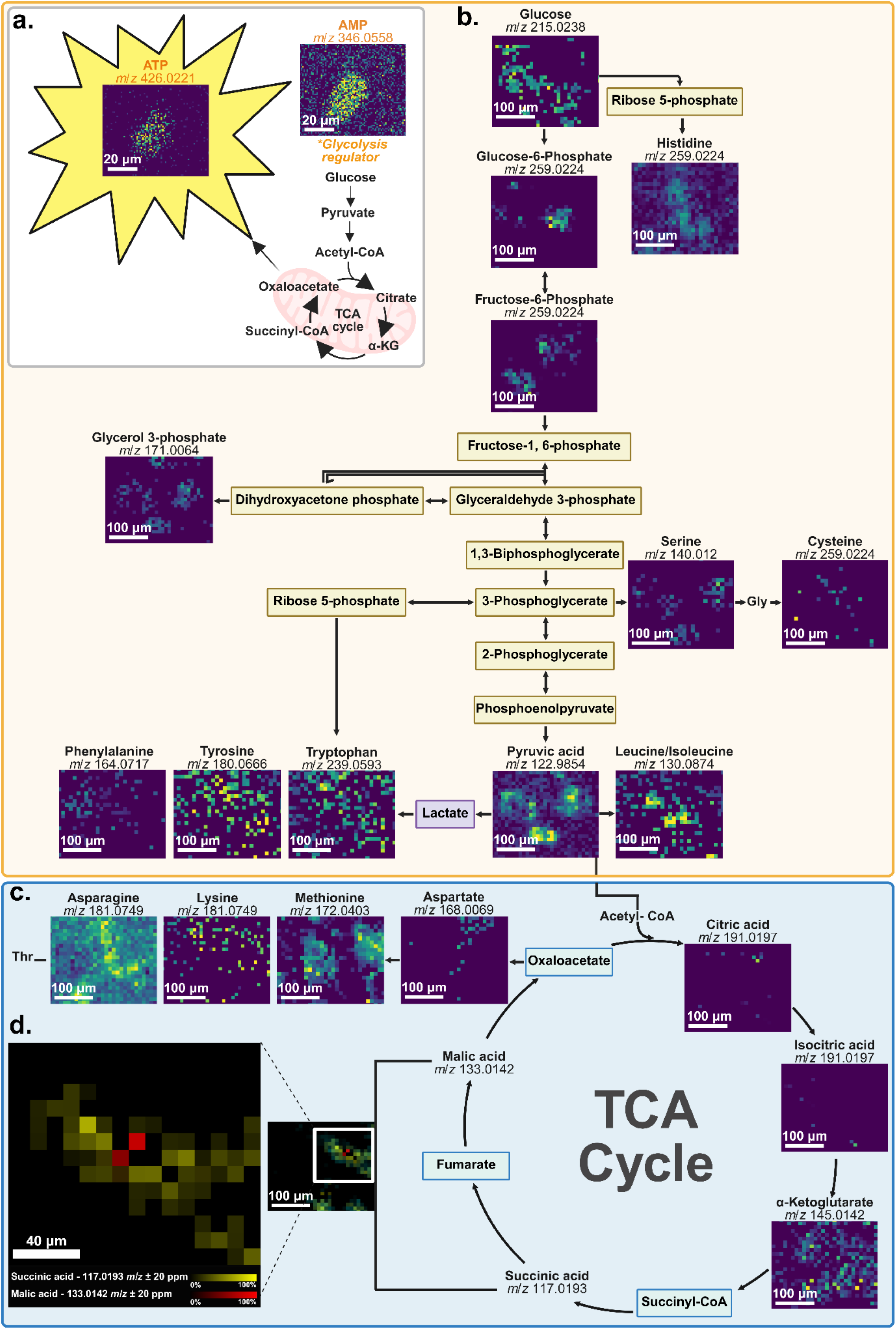
Targeted spatial metabolomics of OVCAR8-RFP cells: glycolysis, the TCA cycle, and amino acid biosynthesis. Targeted spatial metabolomic profiling on OVCAR8-RFP cells over several acquisition runs to investigate metabolites associated with glycolysis, the TCA cycle, and amino acid biosynthesis using MALDI. **(a)** Initial t-MALDI experiments utilizing norharmane matrix enabled the ionization and detection of adenosine triphosphate (ATP) and adenosine monophosphate (AMP). Scale bar, 20 μm. **(b)** TExMS was used to detect multiple metabolic intermediates in glycolysis as well as intermediates linking glycolysis to other pathways. Scale bar, 100 μm. **(c)** Sample preparation and ionization condition optimization were improved metabolite detection, allowing detection of TCA cycle intermediates and amino acid biosynthesis products. Scale bar, 100 μm. **(d)** A representative 2D ion image shows the localization of malic acid (*m/z* 133.0142, red), succinic acid (*m/z* 117.0193, green), and a putative membrane-associated lipid (*m/z* 152.8998, purple). Malic acid was localized within the cell interior and was surrounded by succinic acid, suggestive of the mitochondrial matrix. Scale bar, 40 μm. Data shown in panels **(b–d)** were acquired at a spatial resolution of 10 μm pixel size across multiple acquisitions using different ionization matrices and ion polarities, highlighting that no single analytical approach currently provides comprehensive metabolite coverage. Created in BioRender. Sanchez, L. (2026) https://BioRender.com/ywtalqn

Taking advantage of this improved metabolite coverage of TExMS, we sought to map how OVCAR8- RFP cells undergo metabolic reprogramming allowing them to adapt for rapid proliferation in nutrient-poor conditions such as the tumor microenvironments.^38,39^ This metabolic reprogramming manifests in several downstream effects, including increased glucose uptake, altered gene-driven metabolic regulation, and the diversion of glycolytic and TCA cycle intermediates toward the biosynthesis of key cellular building blocks including amino acids, nucleotides, and lipids.^40,41^ We cultured OVCAR8-RFP cells under serum-treated and serum-starved conditions as a model for nutrient deprivation and observed notable alterations not only in cellular morphology, but also in metabolic profiles. With serum-treated, we observed normal polygonal adherent cell morphology which shifted to a more rounded and non-adherent phenotype in serum-starved cells, consistent with cellular responses (Fig. S8). At the metabolic level, we observed multiple intermediates whose relative abundances were significantly altered under serum-starved conditions (Fig. S9,10) including the key glycolysis regulators pyruvic acid and glucose-6-phosphate (Figure 5). These changes likely reflect stress-associated metabolic adaptations consistent with the metabolic reprogramming observed in cancer cells exposed to nutrient-limited environments.

**Figure 5:**
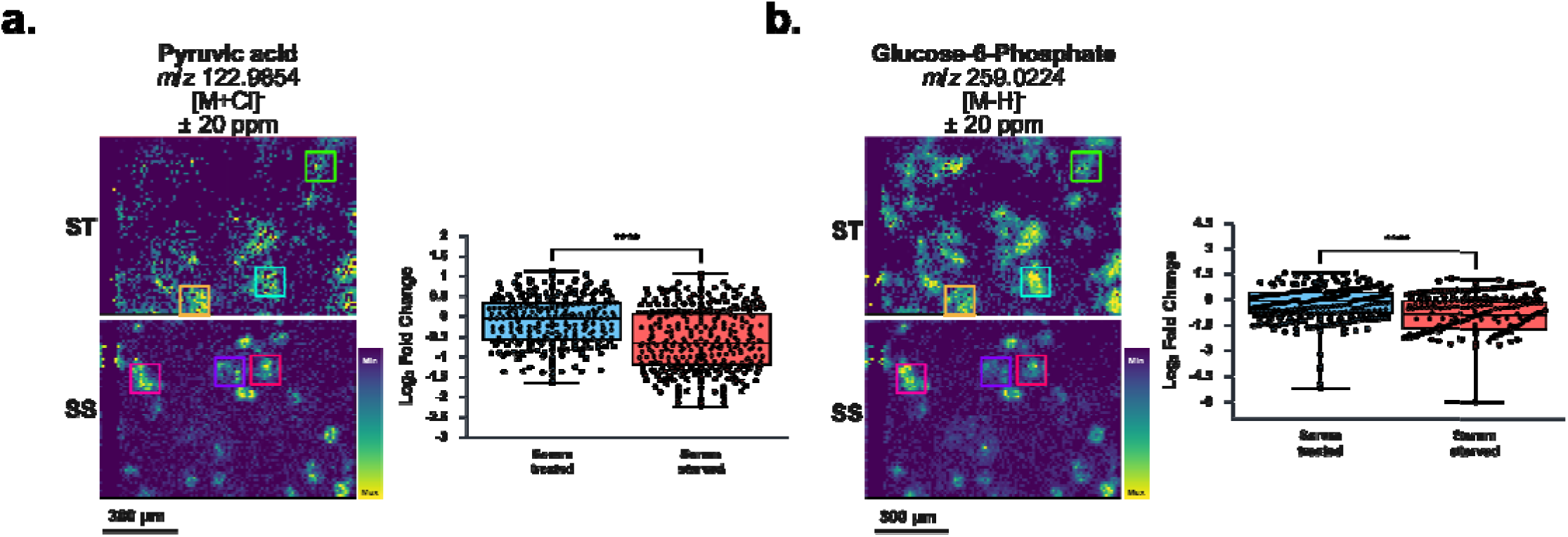
Evaluation of OVCAR8-RFP cells in a nutrient deprived environment. Regions of interest (ROIs) and their corresponding signal intensities were exported for (a) pyruvic acid and (b) glucose-6-phosphate, after which outliers were identified using Grubbs’ test (α = 0.05). A log□ fold-change analysis was subsequently performed on the localized m/z features to compare metabolic alterations between serum-treated (ST) OVCAR8-RFP cells cultured in medium supplemented with 10% fetal bovine serum (FBS) (control) and serum-starved (SS) OVCAR8-RFP cells cultured for 24 hours. Statistical significance was then assessed using Welch’s t-test in BioRender Graphing. The analysis revealed a metabolic flux between the two conditions, reflecting metabolic adaptations of OVCAR8-RFP cells in response to serum starvation. Created in BioRender. Sanchez, L. (2026) https://BioRender.com/1mo26ws

This data further demonstrate that the TExMS workflow can capture spatially resolved metabolic responses associated with altered nutrient availability, energy deficiencies and cellular stress. These findings support the utility of TExMS as a platform for investigating metabolic heterogeneity at the single-cell level.

### Considerations

TExMS can be readily implemented on commercial instruments through modifications to the sample preparation workflow, although several challenges remain. One limitation is throughput, as the complete sample preparation workflow requires approximately four days prior to data acquisition, including extended preparation and incubation times to allow the cells to adhere. Compared with traditional single-cell MALDI-MSI workflows that are 1-2 days in preparation, TExMS requires additional preparation steps, including hydrogel precursor preparation, polymerization, cell seeding, incubation, and sample drying prior to MSI acquisition increasing the overall preparation time despite its relatively rapid data acquisition. Further, we are actively working to improve the observed linear expansion since it does not yet fully reach the theoretical linear expansion predicted by the mechanical stretching geometry. This discrepancy likely reflects current practical constraints associated with OVCAR8-RFP cell adhesion to the hydrogel, such as the effects of tensile expansion on cellular skeleton and cellular elasticity, which we plan to characterize in future studies but should be considered when interpreting the effective spatial expansion achieved by TExMS. Of note for instrumentation considerations is the vacuum stability of hydrogel samples. When mounted using fluorescence-compatible configuration, hydrogels remain structurally stable for only approximately 20 minutes under instrument vacuum before flaking occurs. Consequently, when multimodal imaging is required, the analyzable region was limited to approximately 500 μm^2^ to 1000 mm^2^ due to the restricted acquisition time. To improve stability under vacuum, an alternative mounting strategy was implemented by placing a 18 mm diameter cut of double-sided aluminium tape beneath the desiccated hydrogel. Although this configuration precluded fluorescence imaging, it substantially increased sample stability under vacuum, extending the viable acquisition window from approximately 20 minutes to 1-5 hours and enabling the analysis of larger or multiple regions across the hydrogel. Furthermore, the addition of 100 μm feeler gauge tape enabled fine adjustment of sample height, positioning the hydrogel surface within the optimal focal range of the MALDI laser. These modifications improved laser focusing, ionization efficiency, and signal reproducibility (Figure 3d). Finally, in our metabolic pathway mapping experiments, we conducted several TExMS acquisitions using multiple matrices and in both positive and negative pilates. This highlights that there is currently no single analytical approach that provides comprehensive metabolite coverage over multiple target pathways.

## Conclusions

In conclusion we have demonstrated our TExMS approach using living unfixed cells which relies on native cellular machinery for adherence to the hydrogel surface. The samples are then physically expanded using the iris expansion device to achieve an expansion factor of ∼3.8x on the hydrogel and ∼1.7x for the cells. Our results demonstrate that OVCAR8-RFP epithelial cells adhere to the surface of a functionalized hydrogel and remain structurally intact during mechanical expansion, undergoing expansion without evidence of rupture or membrane failure in response to multiaxial mechanical stretching (Figure 2,4). Furthermore, our findings support that cells desiccation results in measurable metabolites with cellular resolution without negatively affecting ionization efficacy and notably retaining analytes that can be lost with fixation workflows (Figure 3,4). TExMS can be applied broadly to study metabolic heterogeneity at the single-cell level, and we provide a proof of principle study showing the spatial mapping of multiple metabolic intermediates allowing us to characterize the metabolic shifts observed in a representative high grade serous ovarian cancer cell line, OVCAR8, under nutrient deprivation (Figure 5).

Future work will focus on further optimizing the TExMS workflow for broader biological applications in more cell lines with different mutations and different sample types, such as bacteria. We are also furthering developments that will facilitate co-registration of super-resolution tensile expansion microscopy (TExM) optical images^28^ with two-dimensional ion density maps and exploring specific cellular phenomena such as the development of invadopodia on hydrogel surfaces. TExMS results presented here highlight its potential as a platform for studying metabolic heterogeneity across different cell lines at improved spatial resolution without altering the hardware of commercial instruments.

## Methods

The following methods describe the current TExMS workflow. Additional detailed methods for t-MALDI are provided in the Supporting Information (SI).

### Hydrogel precursors stock solutions

To prepare a 2.91% (w/w) sodium alginate stock solution (Alg)(Sigma-Aldrich), 4.5 g of alginic acid sodium salt was gradually added to 150 mL of MilliQ H_2_O (18.2 MΩ·cm) in a glass bottle while initially stirring at 400– 500 rpm. The mixture was then maintained under gentle stirring (50–100 rpm) for two days at room temperature to ensure complete homogeneity and dissolution of the Alg. For the 28.57% (w/w) acrylamide stock (AAm) (Sigma-Aldrich), 20 g of AAm was added to 50 mL of MilliQ H_2_O in a glass bottle and stirred at 300–400 rpm for 1 h, or until fully dissolved. A 1.96% (w/w) N,N′-methylenebisacrylamide (MBA)(Sigma-Aldrich) stock was prepared by dissolving 0.2 g of MBA in 10 mL of MilliQ H_2_O in a 50 mL glass bottle, stirring at 200 rpm for 2 hr. The ammonium persulfate (APS)(Sigma-Aldrich) solution was freshly prepared prior to each hydrogel batch. A 9.1% (w/w) APS solution was made by dissolving 0.2 g of APS in 2 mL of MilliQ H_2_O using a 15 mL conical tube and vortex until completely dissolved.^28^

To prepare approximately 50 mL of hydrogel precursor solution, a sterilized magnetic stir bar is placed into a sterile 100 mL screw-cap glass bottle. A sterile Erlenmeyer flask may also be used, provided it remains sealed throughout the mixing process. Next, 26.63 g of Alg stock solution, 15.56 mL of AAm stock solution, 280.1 μL of MBA, and 171.16 μL of APS were added to yield final concentrations of 1.81% (w/w) Alg, 10.62% (w/w) AAm, with 0.073% (w/w) MBA, and 0.58% (w/w) APS, respectively relative to the mass of AAm. The precursor solution was gently stirred for 45 minutes to ensure complete homogenization. Subsequently, 36.14 μL of N,N,N′,N′-tetramethylethylenediamine (TEMED) is added into the mixture for a final concentration of 0.61% (w/w). The solution was stirred for an additional 15 minutes before being poured into the hydrogel molds for polymerization.The hydrogels were incubated at 35 °C for 5 hours, after which they were removed from the incubator and allowed to continue polymerizing at room temperature.

### Polymerization of hydrogel precursor solution

Following the overnight polymerization at room temperature, the hydrogels were immersed in a freshly prepared 20 mM calcium sulfate dihydrate (CaSO ·2H O)(Sigma-Aldrich) slurry made with MilliQ H_2_O for 12 minutes to complete alginate ionic crosslinking. This step yielded highly stretchable, alginate- Ca^2+^/polyacrylamide (Alg-Ca^2+^/PAAm) double network hydrogels. The hydrogels were then rinsed thoroughly with Milli-Q water and cut into 30 mm diameter plugs (one of three trimmings) using a generic carbon steel leather hole punch purchased from Amazon (ASIN: B0C3LST16X) . Finally, the hydrogel plugs (30 mm) were UV sterilized using a modified 10 L LETORS UV sterilizer (model JY-520, ASIN: B0D9NW6MKC) which has a UV-light band of 220-270 nm, the hydrogels were placed 1 cm away from the UV source for 10 minutes and prepared for subsequent surface functionalization.

### Hydrogel surface functionalization

In a sterile Petri dish (60 mm), hydrogels were centered using sterile tweezers. The 30 mm hydrogels were coated with 1 mL of 0.1 mg/mL PDL and incubated for 1 h at room temperature. Excess PDL was removed by rinsing the hydrogels three times with 1 mL sterile MilliQ H_2_O. PDL-coated hydrogels were then air-dried for 15–30 minutes under ambient conditions, until the surface appeared visibly dry. Initial screening experiments (Fig. S11) utilized a simple surface treatment approach in which poly-D-lysine (PDL) was diffused onto and rehydrated within the hydrogel surface.^42^ While the PDL treatment allowed for minimal cell adhesion to the hydrogel substrate, the cells did not withstand mechanical stress testing. A 300 μL aliquot of a 500 μM sulfo-SANPAH diluted in 100 mM HEPES buffer was added to fully cover the surface. The hydrogels were then placed inside the LETORS UV sterilizer, positioned approximately 1 cm from the UV light source (220–270 nm), and irradiated for 15 minutes. The sulfo-SANPAH activation procedure is repeated twice for each hydrogel to ensure uniform surface coverage and effective functionalization. Following each UV exposure, the hydrogels were rinsed three times with 1 mL of 100 mM HEPES buffer to remove unreacted sulfo-SANPAH. After the final HEPES rinse, 400 μL of 350 μg/mL human plasma FN (Thermo Fisher) diluted in MilliQ H_2_O was added to the surface of each functionalized hydrogel. The protein-coated hydrogels were then incubated at 4°C in a humidified environment for 24 hours. Following the 24 hours fibronectin (FN) incubation, the hydrogels increased in diameter by an additional 2–5 mm and were trimmed a second time (two of three trimmings) to a diameter of 30 mm to fit within 6-well plates. Followed by UV sterilization for 15 minutes and subsequently rinsed three times with 1mL 50 mM HEPES buffer to remove excess FN. The hydrogels were then transferred from the Petri dish to a sterile 6-well plate (Falcon® clear) for cell seeding.

### Cell culture

OVCAR8-RFP^29^ cells were maintained in Dulbecco’s Modified Eagle Medium (DMEM (1×)) (GenClone®) containing high glucose, L-glutamine, and sodium pyruvate, supplemented with 10% fetal bovine serum (FBS) (Sigma-Aldrich) and 1% penicillin–streptomycin (PenStrep)(Gibco). Prior to passaging, cells were rinsed with 1x phosphate buffer saline (1x PBS) (GenClone®), then detached and resuspended using 0.25% trypsin– EDTA with phenol red (TR-EDTA) (Gibco). Cells were passed every 3–4 days and maintained at 37 °C in a humidified incubator with 5% CO_2_. All experiments were performed using cells passaged 20 - 40. Cells exceeding passage 40 were discarded and replaced.

### Fixation

Chemical fixation was performed using a stock 30% formaldehyde solution (HCOH)(Roth) containing low methanol, which was diluted in 1× PBS to obtain a final concentration of 4% HCOH. Cells were coated with 500 μL of the 4% HCOH solution for 5 minutes. Following fixation, the solution was removed, and the cells were washed and dried according to the procedures previously reported.^12^

### t-MALDI sample preparation

#### Cell seeding

In a Millicell EZ Slide 8-well chamber (Sigma-Aldrich), 500 µL of 0.1 mg/mL PDL was added to each of the eight wells and allowed to incubate undisturbed at room temperature for 1 hour to functionalize the glass substrate. Following incubation, excess PDL was removed following rinsing the wells three times with sterile MilliQ H_2_O. The slides were then air-dried for 15 minutes under ambient conditions prior to cell seeding.

OVCAR8 cells were resuspended and counted. Cells were mixed with trypan blue at a 1:1 (v/v) ratio to assess viability and determine cell concentration. Cells were then diluted to a final concentration of 10,000 viable cells per well and seeded onto the PDL-functionalized slides for a final volume of 500μL of DMEM (1x) and incubated for 24 hours at 37 °C and 5% CO_2_.

#### Fixation & staining

After the fixation steps previously mentioned above, a working solution of Hoechst 33342 trihydrochloride trihydrate (H3570; Thermo Fisher Scientific) was prepared by diluting a 16.2 mM stock solution to 16.2 µM (1μg/mL) in 1× PBS. A volume of 250 µL of the Hoechst nuclear stain was added to wells containing fixed OVCAR8 cells and incubated for 5 minutes at room temperature, protected from light. Following incubation, samples were rinsed three times with 500 µL of 1× PBS. 250 µL of CellMask™ Green Actin Tracking Stain diluted in (1μg/mL) 1x PBS was added to the fixed cells and incubated for 30 minutes at room temperature, protected from light. After staining, samples were rinsed three times with 1 mL of 150 mM ammonium acetate (NH_4_OAc)(Thermo Fisher) to remove excess dye and to prepare the samples for downstream mass spectrometry–compatible workflows. Fluorescence images were acquired using an Olympus VS200 Research Slide Scanner configured for fluorescence, brightfield, and darkfield imaging. Image acquisition and processing were performed using OlyVIA software (Evident) with an X-Cite NOVEM light source and a Hamamatsu

ORCA-Fusion camera.

#### Matrix deposition

Norharmane (NH) (Sigma-Aldrich) matrix was applied via sublimation for high resolution imaging (1-5 μm). The NH matrix was prepared by dissolving norharmane at 5 mg/mL in 100% ethanol (EtOH)(Roth).

Sublimation was performed using a Bruker superlimator with a preheating temperature of 140 °C, a cooling temperature of 10 °C, and a sublimation temperature of 160 °C for a total of 500 seconds and subjected to recrystallization at 65 °C for 2 minutes and 150 seconds using a makeshift vapor chamber constructed from a large petri dish and a filter paper soaked with 1 mL of 0.5% EtOH prepared in MilliQ H_2_O.

#### Ion optics parameters

Single-cell MSI data were acquired at a (pixel size 1 μm) using a Bruker timsTOF mass spectrometer modified for t-MALDI-MSI, as described in detail in.^1^ The MS1 mass range was set to 5–1000 *m/z*. Ion optics settings for MS1 acquisition in negative-ion mode were as follows: the deflection delta was set to −70.0 V, and the MALDI plate offset was maintained at 50.0 V. Funnel 1 RF, Funnel 2 RF, and Multipole RF were each operated at 400.0 Vpp. The quadrupole low-mass cutoff was set to 50 m/z with an ion energy of 5.0 eV.

Collision cell parameters included a collision energy of 10 eV and a collision RF of 720.0 Vpp, with a transfer time of 60 μs and a pre-pulse storage time of 5.0 μs. High-sensitivity and focus-mode settings were disabled. Laser parameters were configured as follows: Burst = 1, number of shots: 15, frequency: 10,000 Hz, laser power: 80%, and a custom application mode.

### MALDI-2 (microgrid) sample preparation

Sample preparation for cell seeding and fixation followed previously described t-MALDI preparation steps.

#### Matrix deposition

N-(1-naphthyl)ethylenediamine dihydrochloride (NEDC)(Sigma-Aldrich) matrix solution was prepared at a concentration of 3 mg/mL in 100% MeOH. Using a Bruker Superlimator, 1 mL of the NEDC solution was added to the sublimation crucible. The system was programmed with a preheating temperature of 85 °C, a cooling temperature of 0 °C, and a sublimation temperature of 160 °C, with a total sublimation duration of 180 seconds. The samples were then subjected to recrystallization at 65 °C for 2 minutes and 150 seconds using a makeshift vapor chamber constructed as previously mentioned in the t-MALDI sample preparation section.

#### Ion optics parameters

Single-cell MSI data (pixel size 5 μm) were acquired on a Bruker timsTOF fleX MALDI-2 instrument in negative-ion polarity. The MS1 mass range was set to 50–650 *m/z*. Ion optics settings for MS1 acquisition in negative mode were as follows: the deflection delta was −70.0 V, and the MALDI plate offset was 50.0 V. Funnel 1 RF, Funnel 2 RF, and Multipole RF were set to 125.0 Vpp, 200.0 Vpp, and 200.0 Vpp. The quadrupole low-mass cutoff was maintained at 50 m/z with an ion energy of 5.0 eV. Collision cell parameters included a collision energy of 10 eV and a collision RF of 350.0 Vpp, with a transfer time of 55 μs and a pre- pulse storage time of 5.0 μs. High-sensitivity and focus-mode settings were disabled. Laser parameters were configured as follows: Burst = 1, number of shots: 25, frequency: 10,000 Hz, laser power: 80%, with the global attenuator was set to an additional 10%., with a custom application mode using SmartBeam enabled. The scan range in both X and Y was set to 1 μm, yielding a final field size in both X and Y of 5 μm.

### TExMS sample preparation

#### Cell seeding

OVCAR-RFP cells were seeded at high confluence directly onto the center of the hydrogel and allowed to attach for 45 minutes at 37 °C with 5% CO_2_ before gently adding 7 mL of DMEM (1×) medium. The samples were then incubated for an additional 24 hours to allow for adhesion and proliferation. Following the 24 hour of incubation, cells were assessed for cell morphology and substrate adhesion. DMEM (1×) was removed, and wells were rinsed three times with 1× PBS to remove residual serum components. A volume of 1 mL of 8.1 µM Hoechst 33342 solution was added to each well to ensure complete coverage of the cell monolayer. Samples were protected from light and incubated for 10 minutes at 37 °C and 5% CO_2_ Following incubation, excess stain was removed, and cells were rinsed three times with 2 mL of 1× PBS to eliminate unbound dye. The cell monolayer was subsequently rehydrated with fresh DMEM (1×) and returned to the incubator at 37 °C and 5% CO_2_ until proceeding with mechanically induced stretching. Fluorescence images of pre- and post-expanded cells were acquired using a Zeiss Axio Imager Z1 widefield microscope equipped with a 2.5×/0.12 NA objective (working distance = 8.7 mm). For the pre-expanded condition, hydrogel samples were trimmed to a 15 mm diameter to fit onto a glass microscope slide. For post-expanded samples, hydrogels were trimmed to 30 mm diameter to fit onto the iris mounting apparatus and subjected to a maximum multiaxial expansion of ∼3.8x, followed by desiccation at 32 °C for 5 hours, desiccation is a critical step for eventual MSI, thus we sought to test dessicated samples early for subsequent MSI compatibility. Once desiccated 18 mm coverslip was placed beneath the sample before desiccation to provide an optically transparent support for the hydrogel during desiccation and enable fluorescence imaging (data not included). To further evaluate expansion effects on cellular morphology, the experiment was repeated under similar conditions but without the desiccation step after expansion. Static fluorescence images were analyzed using ImageJ (fiji). Cell area measurements were performed manually. In contrast, nucleus area measurements were obtained automatically using the particle analysis function.

#### Matrix deposition

For high-resolution MALDI-MSI, matrix crystal size should be smaller than the acquisition pixel size to ensure efficient laser energy absorption by the matrix and effective energy transfer to analyte molecules. To improve matrix homogeneity, multiple deposition conditions were evaluated, with the optimization process described in the Supporting Information (Table S11). α-Cyano-4-hydroxycinnamic acid (CHCA) (Sigma-Aldrich) was recrystallized in-house prior to use. CHCA & NEDC was prepared at a concentration of 5 mg/mL in a solvent mixture consisting of 9:1 (v/v) acetonitrile (ACN) (Sigma-Aldrich) and MilliQ H_2_O containing 0.1% trifluoroacetic acid (TFA)(Sigma-Aldrich) only for the CHCA matrix. 2D matrix deposition was performed using an HTX TM-Sprayer™ (HTX Technologies LLC) with nitrogen gas at a flow rate of 3 L/min (10 psi). Sprayer parameters were optimized as follows: nozzle temperature set to 90 °C, 12 passes in a crisscross pattern, solvent composition of 50:50 (v/v) HPLC-grade MeOH and MilliQ H_2_O, flow rate of 0.2 mL/min, track spacing of 3 mm, and nozzle height of 40 mm above the sample surface. The use of a high organic solvent content and elevated nozzle temperature promoted the formation of small, uniform CHCA crystals, enhancing matrix homogeneity. Similarly to the CHCA sprayer parameters NEDC matrix deposition was applied at 60 °C to avoid clogging the nozzle. These conditions produced a homogeneous matrix coating, enabling improved ionization and allowing putative metabolite annotations from the OVCAR8-RFP in-house metabolite library (Fig. S12-Fig. S14).

#### Ion optic parameters

Single-cell MSI data (pixel size 10 μm) were acquired in both positive and negative-ion modes using a Bruker timsTOF fleX mass spectrometer. The MS1 mass range was set to 50–1200 m/z. The deflection delta was set to 75.0 V, and the MALDI plate offset was also maintained at 75.0 V. Peak-to-peak voltages for the ion optics were configured as follows: Funnel 1 RF at 150.0 Vpp, Funnel 2 RF at 200.0 Vpp, and Multipole RF at 200.0 Vpp. The quadrupole low-mass cutoff was set to 50 m/z with an ion energy of 5.0 eV. Collision cell parameters included a collision energy of 10 eV and a collision RF of 350.0 Vpp, with a transfer time of 55 μs and a pre- pulse storage time of 5.0 μs. High-sensitivity and focus-mode settings were disabled. Laser parameters were configured as follows: Burst = 1, number of shots: 1000, frequency: 5,000 Hz, laser power: 70%, and a, and a custom application mode with SmartBeam enabled. The scan range in both X and Y was set to 6 μm, producing a final field size in both X and Y of 10 μm. MSI data were processed and visualized using SCiLS Lab Pro (Bruker). All ion images were normalized to the total ion count (TIC), and no denoising algorithms were applied during data processing.

### Iris control

Pre-expansion hydrogel samples containing the cells are trimmed for a third time (three out of three trimmings) to a final diameter of 30 mm, allowing it to fit onto the 3D printed mounting apparatus that will enable it to be loaded onto the iris expansion device (Fig. S15). The expansion parameters such as speed and expansion factor can be precisely controlled through a custom built PCB-based control box containing a script written in C++ and run via Arduino software as previously described with higher detail.^28^ Once the hydrogel is mounted and secured into place, the mounting apparatus was disengaged and the system was set to the slowest expansion speed of 0.01 cm/s and a maximum multiaxial expansion factor of ∼3.8x.

### Incubation and drying

Falcon® clear 6-well plates containing the hydrogels were incubated at 37 °C in a humidified atmosphere with 5% CO_2_ for 24 hours, using a hydration plate to maintain moisture. Following incubation, the hydrogels were mechanically expanded using the iris expansion device. The expanded hydrogels were secured with magnetic drying rings, where excess hydrogels hanging from the sides of the clips were trimmed using a box cutter, 18 mm (0.13-0.17 mm) coverslip was added to the underside of the sample before drying. Samples were dried at ambient conditions for 4-6 hours while continuously rotated on a custom-built spinner.^43^

### In-house OVCAR8-RFP metabolome library

LC-MS separation for the preparation of the OVCAR8-RFP metabolome library, OVCAR8-RFP cells were incubated at 37°C with 5.0% CO_2_ in a T225 flask until reaching approximately 100% confluency. Once the appropriate confluency was achieved, the cells were detached, resuspended, and counted using ACCURIS instruments Quadcount with trypan blue at a 1:1 (v/v) ratio to obtain a final concentration of 6 million cells. The cell suspension was transferred into a 5 mL conical tube and centrifuged at 900 rpm for 5 minutes. The supernatant was carefully removed without disturbing the pellet, which was then resuspended in 1 mL of 1× PBS and centrifuged at 900 rpm for 2 minutes. This washing step was repeated three times. After the final wash, the cell pellet was stored at −80 °C in preparation for lyophilization. Following lyophilization, the dried cell mass was gently broken apart using a stainless-steel spatula and processed for solvent-based extraction. A 1:1 (v/v) mixture of ethyl acetate (EtOAc)(Sigma-Aldrich) and MeOH was prepared, and 1 mL of the EtOAc:MeOH solution was added to each lyophilized sample. The samples were sonicated for 1 h in an ice- water bath to prevent overheating the samples. For LC-MS/MS analysis, the extracted samples were centrifuged at 13,500 rpm for 10 minutes to remove large particulates. The supernatant was carefully transferred to 4 mL glass pre-weighed vials, ensuring that the remaining cell debris was not disturbed. Samples were then prepared for LC–MS/MS at a final concentration of 1 mg/mL. Data acquisition was performed on a Bruker timsTOF fleX mass spectrometer coupled to a Bruker Elute UHPLC HPG 1300 system. Analyses were carried out using both reverse-phase liquid chromatography (RPLC) and hydrophilic interaction liquid chromatography (HILIC) in positive and negative ion polarity modes. For RPLC a volume of crude extract was injected onto a Kinetix C18 column (1.7 μm, 50 × 2.1 mm, 100 Å; P/N: 00B-4475-AN). Mobile phase A1 consisted of LC-grade H_2_O with 0.1% optima grade formic acid (FA) (Thermo Fisher), and mobile phase B1 consisted of Optima LC-MS grade ACN (Sigma-Aldrich) with 0.1% FA. Chromatographic separation was achieved using a 15-minute gradient at a constant flow rate of 0.5 mL/min. The gradient profile was as follows: 0.0–0.5 minutes, 95% A1 (isocratic hold); 0.5–9.5 minutes, linear gradient to 100% B1; 9.5–11.5 minutes, column wash at 100% B1; 11.5–13.0 minutes, return to 95% A1; 13.0–15.0 minutes, re-equilibration at 95% A1. For HILIC, an aliquot of crude extract was injected onto an InfinityLab Poroshell HILIC column (2.1 × 50 mm, 1.9 μm; P/N: 699675-901). Mobile phase A2 consisted of 10 mM optima LC-MS grade ammonium acetate (NH_4_OAc)(Thermo Fisher) with optima LC-MS grade acetic acid (AcOH) (Thermo Fisher), and mobile phase B2 consisted of ACN:100 mM NH_4_OAc (90:10)with AcOH. Separation was performed using a 16-minute gradient at a constant flow rate of 0.2 mL/min. The gradient program was as follows: 0.0–3.50 minutes, 95% B2 (isocratic hold); 3.50–4.00 minutes, linear gradient; 4.00–6.00 minutes, 80% B2 (isocratic hold); 6.00–11.50 minutes, linear gradient to 50% B2; 11.50–13.50 minutes, 50% B2 (isocratic hold); 13.50–15.00 minutes, return to 95% B2; and 15.00–16.00 minutes, re-equilibration at 95% B2. Data acquisition for both MS^1^ and MS^2^ was performed on a Bruker timsTOF fleX system, using comparable instrument parameters across positive- and negative-ion polarities for both HILIC and RPLC analyses. For MS1 acquisition in both negative- and positive-ion polarities, the mass range was set to 20–2000 m/z with a spectral acquisition rate of 6.00 Hz. Electrospray ionization (ESI) source parameters were configured as follows: endplate offset at 500 V, capillary voltage at 4200 V, nebulizer pressure at 2.8 bar, dry gas flow at 10.0 L/min, and drying temperature at 230 °C. Ion-optics settings were adjusted for each polarity. The deflection 1 delta was set to −60.0 V in negative-ion mode and +60.0 V in positive-ion mode. For both modes, Funnel 1 RF and Funnel 2 RF were set to 200.0 Vpp, and the multipole RF was maintained at 300.0 Vpp. The quadrupole low-mass cutoff was set to 20.0 m/z with an ion energy of 5.0 eV with a pre-pulse storage of 5.0 μs. Collision cell settings included a collision energy of 10.0 eV for negative-ion mode and 7.0 eV for positive-ion mode. For both modes, the collision RF ranged from 200.0 Vpp to 700.0 Vpp. Transfer times ranged from 13.5 μs to 54.0 μs in negative-ion mode, and from 20.0 μs to 80.0 μs in positive-ion mode. For MS^2^ acquisition, parameters for both positive- and negative-ion modes were configured as follows. The mass range was set to 100–2000 m/z with a spectral acquisition rate of 10.00 Hz. ESI source parameters were identical to those used for MS1, with the exception of the capillary voltage, which was increased to 4500 V. Ion-optics settings were the same as in MS^1^, except that the quadrupole low-mass cutoff was raised to 100 m/z while maintaining an ion energy of 5.0 eV. Collision cell energies were set to 10.0 eV for negative-ion mode and 30.0 eV for positive-ion mode. For both polarities, the collision RF was maintained at 700.0 Vpp, with a pre-pulse storage time of 5.0 μs. Transfer times were 46.0 μs in negative-ion mode and 80.0 μs in positive-ion mode. MS^2^ acquisition parameters were configured to target the top ten precursor peaks with an absolute intensity threshold of 350 counts. Automatic MS^2^ CID settings were used, with MS^2^ spectral acquisition constrained to a minimum of 30.00 Hz and a maximum of 16.00 Hz, a fixed MS^2^ acquisition rate of 10.00 Hz, and a target intensity of 20,000. All MS^1^ and MS^2^ raw files were analyzed using MetaboScape (Bruker). Further dereplication and biological annotation were performed using Global GNPS2^33^ and HMDB.^34^

### Statistical analysis

Static fluorescence microscopy images (.czi files) were processed using Image J (fiji). Images were first set to scale using the rectangle selection tool and the embedded scale information to convert pixel measurements into calibrated distances and areas of measurements. The images were then converted from RGB to 8-bit grayscale, and intensity thresholds were adjusted to distinguish cells and nuclei from the background.

Background artifacts and small particles were further reduced using the erode and dilate functions before analysis. Whole-cell boundaries could not be reliably segmented using automated image analysis; therefore, cell area measurements were collected manually. In contrast, nuclei area measurements were acquired automatically using the Analysis Particles function after thresholding., with the particle size threshold set from 0 to infinity. Regions of interest (ROIs) were automatically outlined and quantified by the software. Area measurements were exported from ImageJ (fiji), organized in Microsoft Excel, and subsequently imported into BioRender for graphing and statistical analysis. Cell and nuclear area measurements obtained from matched pre- and post-expansion samples were analyzed using a one tailed paired t-test under parametric data distribution.

## Supporting information

Supplemental Information OVCAR8 Measurement Summary

## Acknowledgements

This research was financially supported by an Allen Distinguished Investigator Award supported by Allen Family Philanthropies, a Scialog program sponsored jointly by Research Corporation for Science Advancement, Activating Innovative Graduate Research Award and the Gordon and Betty Moore Foundation and includes grant numbers 28410 (LK) and 28411 (LMS), NIH NIGMS R35GM142466 (LK), NIH NIGMS K12GM139185 (EAO), and NIH NICHD T32HD108079 (EAO). We also thank members of the Andresen Eguiluz, and Kisley labs for useful discussions, and the entire Sanchez lab for discussion. We thank Bruker Daltonics (Bremen, Germany) for support of the project through the RCMSI in Münster.

