## Supplemental Information for "Tensile Expansion Mass Spectrometry for single cell metabolomics imaging"

**Table S1:** In-house metabolome library for OVCAR8-RFP cells.

| Name | Annotation source | Molecular formula | Exact mass | M+H | M-H | M+Cl |
|---|---|---|---|---|---|---|
| a-Ketoglutarate | HMDB | $C_5H_6O_5$ | 146.021525 | 147.02935 | 145.0137 | 180.990378 |
| Adenine | HMDB | $C_5H_5N_5$ | 135.054495 | 136.06232 | 134.04667 | 170.023348 |
| Adenonsine | Metaboscape | $C_{10}H_{13}N_5O_4$ | 267.096755 | 268.10458 | 266.08893 | 302.065608 |
| ADP | HMDB | $C_{10}H_{15}N_5O_{10}P_2$ | 427.029421 | 428.037246 | 426.021596 | 461.998274 |
| a-Ketoglutarate | HMDB | $C_5H_6O_5$ | 146.021525 | 147.02935 | 145.0137 | 180.990378 |
| AMP | HMDB | $C_{10}H_{14}N_5O_7P$ | 347.063088 | 348.070913 | 346.055263 | 382.031941 |
| Arginine | Metaboscape | $C_6H_{14}N_4O_2$ | 174.111676 | 175.119501 | 173.103851 | 209.080529 |
| Asparagine | Metaboscape | $C_4H_8N_2O_3$ | 132.053493 | 133.061318 | 131.045668 | 167.022346 |
| Aspartate | Metaboscape | $C_4H_7NO_4$ | 132.029684 | 133.037509 | 131.021859 | 166.998537 |
| Aspartic acid | Metaboscape | $C_4H_7NO_4$ | 133.037509 | 134.045334 | 132.029684 | 168.006362 |
| ATP | HMDB | $C_{10}H_{16}N_5O_{13}P_3$ | 506.995754 | 508.003579 | 505.987929 | 541.964607 |
| ATP | HMDB | $C_{10}H_{16}N_5O_{13}P_3$ | 506.995754 | 508.003579 | 505.987929 | 541.964607 |
| Carnitine | Metaboscape | $C_7H_{15}NO_3$ | 161.105193 | 162.113018 | 160.097368 | 196.074046 |
| Cholesterol | HMDB | $C_{27}H_{46}O$ | 386.354865 | 387.36269 | 385.34704 | 421.323718 |
| Choline | Metaboscape | $C_5H_{14}NO$ | 104.107539 | 105.115364 | 103.099714 | 139.076392 |
| Citric acid | HMDB | $C_6H_8O_7$ | 192.027005 | 193.03483 | 191.01918 | 226.995858 |
| Citrulline | Metaboscape | $C_6H_{13}N_3O_3$ | 175.095691 | 176.103516 | 174.087866 | 210.064544 |
| Creatine | Metaboscape | $C_4H_9N_3O_2$ | 131.069477 | 132.077302 | 130.061652 | 166.03833 |
| Creatine phosphate | Metaboscape | $C_4H_{10}N_3O_5P$ | 211.03581 | 212.043635 | 210.027985 | 246.004663 |
| Cystiene | Metaboscape | $C_3H_7NO_2S$ | 121.019751 | 122.027576 | 120.011926 | 155.988604 |
| Folic acid | GNPS2 | $C_{19}H_{19}N_7O_6$ | 441.139683 | 442.147508 | 440.131858 | 476.108536 |
| Frutose-6-Phospate | HMDB | $C_6H_{13}O_9P$ | 260.029723 | 261.037548 | 259.021898 | 294.998576 |
| Glucose | GNPS2 | $C_6H_{12}O_6$ | 180.06339 | 181.071215 | 179.055565 | 215.032243 |
| Glucose-6-Phosphate | HMDB | $C_6H_{13}O_9P$ | 260.029723 | 261.037548 | 259.021898 | 294.998576 |
| Glutamate | Metaboscape | $C_5H_8NO_4$ | 146.045334 | 147.053159 | 145.037509 | 181.014187 |
| Glutamine | Metaboscape | $C_5H_{10}N_2O_3$ | 146.069143 | 147.076968 | 145.061318 | 181.037996 |
| Glutathione | Metaboscape | $C_{10}H_{17}N_3O_6S$ | 307.083809 | 308.091634 | 306.075984 | 342.052662 |
| Glycerol 3-phosphate | HMDB | $C_3H_9O_6P$ | 172.013678 | 173.021503 | 171.005853 | 206.982531 |
| Glycerophosphocholine | Metaboscape | $C_8H_{20}NO_6P$ | 257.102827 | 258.110652 | 256.095002 | 292.07168 |
| Guanosine | Metaboscape | $C_{10}H_{13}N_5O_5$ | 283.09167 | 284.099495 | 282.083845 | 318.060523 |
| Histidine | Metaboscape | $C_6H_9N_3O_2$ | 155.069477 | 156.077302 | 154.061652 | 190.03833 |
| Hypoxanthine | Metaboscape | $C_5H_4N_4O$ | 136.038511 | 137.046336 | 135.030686 | 171.007364 |

| | | | | | | |
|---|---|---|---|---|---|---|
| Indolin | Metaboscape | $C_8H_9N$ | 119.073499 | 120.081324 | 118.065674 | 154.042352 |
| Citric/isocitric acid | HMDB | $C_6H_8O_7$ | 192.027005 | 193.03483 | 191.01918 | 226.995858 |
| Kynurenic acid | HMDB | $C_{10}H_7NO_3$ | 189.042594 | 190.050419 | 188.034769 | 224.011447 |
| Lactate | HMDB | $C_3H_5O_3$ | 89.02387 | 90.031695 | 88.016045 | 123.992723 |
| Lactic acid | HMDB | $C_3H_6O_3$ | 90.031695 | 91.03952 | 89.02387 | 125.000548 |
| Leucine/isoleucine | Metaboscape | $C_6H_{13}NO_2$ | 131.094629 | 132.102454 | 130.086804 | 166.063482 |
| Linoleic acid | HMDB | $C_{18}H_{32}O_2$ | 280.24023 | 281.248055 | 279.232405 | 315.209083 |
| Lysine | Metaboscape | $C_6H_{14}N_2O_2$ | 146.105528 | 147.113353 | 145.097703 | 181.074381 |
| Malic acid | HMDB | $C_4H_6O_5$ | 134.021525 | 135.02935 | 133.0137 | 168.990378 |
| Methionine | Metaboscape | $C_5H_{11}NO_2S$ | 149.051051 | 150.058876 | 148.043226 | 184.019904 |
| Orotic acid | HMDB | $C_5H_4N_2O_4$ | 156.017108 | 157.024933 | 155.009283 | 190.985961 |
| Palmitic acid | HMDB | $C_{16}H_{32}O_2$ | 256.24023 | 257.248055 | 255.232405 | 291.209083 |
| Penicillin G | Metaboscape | $C_{16}H_{18}N_2O_4S$ | 334.09873 | 335.106555 | 333.090905 | 369.067583 |
| Phenol red | Metaboscape | $C_{19}H_{14}O_5S$ | 354.056197 | 355.064022 | 353.048372 | 389.02505 |
| Phenyanalanine | Metaboscape | $C_9H_{11}NO_2$ | 165.078979 | 166.086804 | 164.071154 | 200.047832 |
| Pyruvate | HMDB | $C_3H_3O_3$ | 87.00822 | 88.016045 | 86.000395 | 121.977073 |
| Pyruvic acid | HMDB | $C_3H_4O_3$ | 88.016045 | 89.02387 | 87.00822 | 122.984898 |
| Serine | Metaboscape | $C_3H_7NO_3$ | 105.042594 | 106.050419 | 104.034769 | 140.011447 |
| Succinic acid | HMDB | $C_4H_6O_4$ | 118.02661 | 119.034435 | 117.018785 | 152.995463 |
| Succinic anhydride | HMDB | $C_4H_4O_3$ | 100.016045 | 101.02387 | 99.00822 | 134.984898 |
| Taurine | Metaboscape | $C_2H_7NO_3S$ | 125.014666 | 126.022491 | 124.006841 | 159.983519 |
| Theronine | Metaboscape | $C_4H_9NO_3$ | 119.058244 | 120.066069 | 118.050419 | 154.027097 |
| Tryptophan | Metaboscape | $C_{11}H_{12}N_2O_2$ | 204.089878 | 205.097703 | 203.082053 | 239.058731 |
| Tyrosine | Metaboscape | $C_9H_{11}NO_3$ | 181.073894 | 182.081719 | 180.066069 | 216.042747 |
| Uracil | Metaboscape | $C_4H_4N_2O_2$ | 112.027278 | 113.035103 | 111.019453 | 146.996131 |
| Uridine | GNPS2 | $C_9H_{12}N_2O_6$ | 244.069538 | 245.077363 | 243.061713 | 279.038391 |
| Valine | HMDB | $C_5H_{11}NO_2$ | 117.078979 | 118.086804 | 116.071154 | 152.047832 |
| Xanthine | Metaboscape | $C_5H_4N_4O_2$ | 152.033426 | 153.041251 | 151.025601 | 187.002279 |

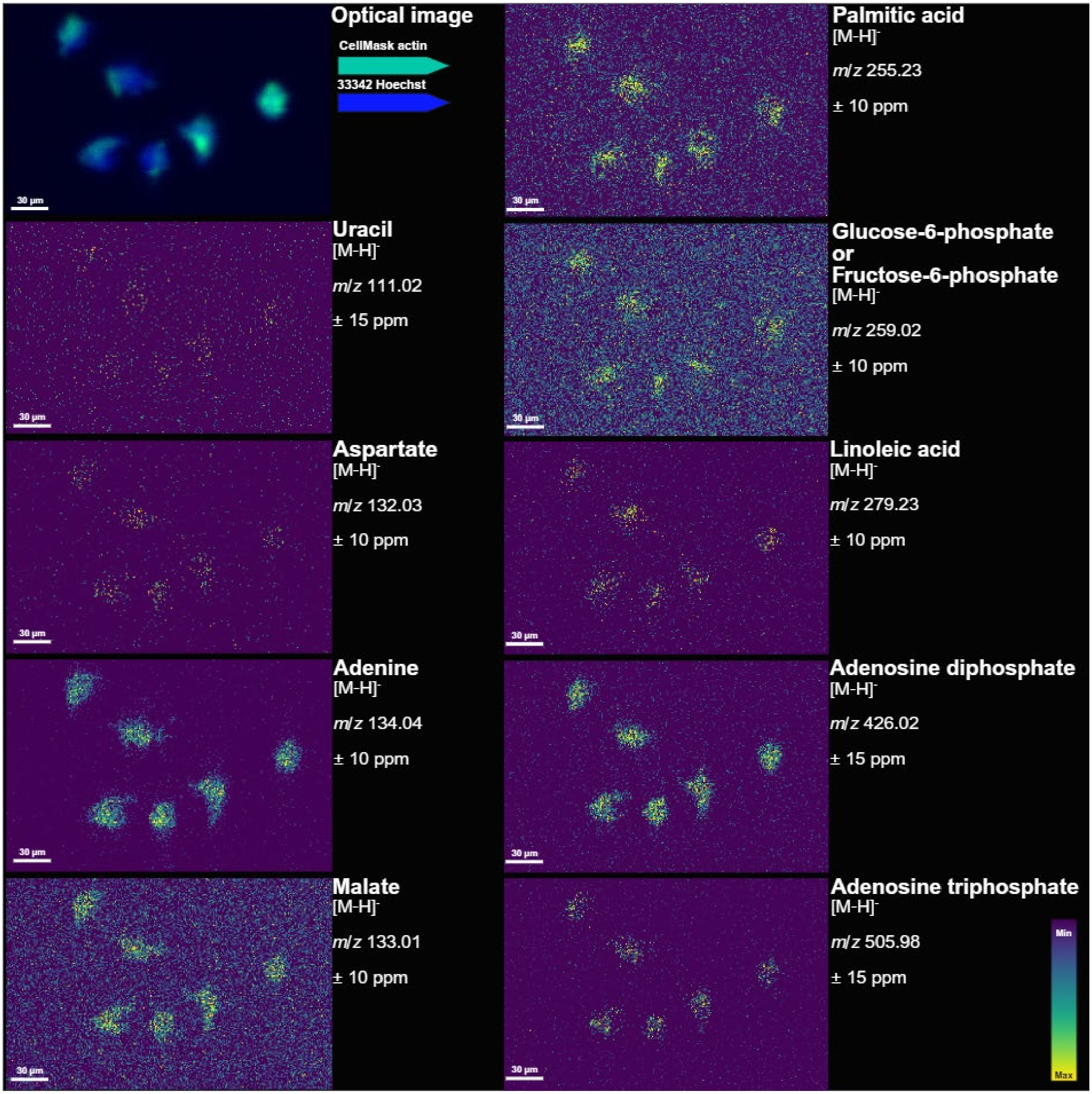

**Figure S1: *t-MALDI data*.** In-house built transmission mode-MALDI2 prototype allowed for high resolution imaging on fixed OVCAR8 cells. 2D ion images with their respective annotations.

**Table S2:** Sublimation parameters for 1 μm–resolution data collection

| | |
|---|---|
| **Matrix (concentration & solvent)** | Norharmane ( 5mg/mL dissolved in 100% EtOH) |
| **Preheating temperature (°C)** | 140 |
| **Sublimation temperature (°C)** | 160 |
| **Cooling temperature (°C)** | 10 |
| **Sublimation time (s)** | 500 |
| **Matrix thickness (mg)** | 2 |

**Table S3:** MS ion optics for t-MALDI-MSI 1µm data. In-house prototype built on a Bruker timsTOF flex.

| **MS Setting** | | **Tune** | |
|---|---|---|---|
| Scan begins (m/z) | 50 | MALDI Plate offset (V) | 50.0 |
| Scan ends (m/z) | 1000 | Deflection 1 delta (V) | -70.0 |
| Ion polarity | Negative | Funnel 1 RF (Vpp) | 400.0 |
| Scan mode | MS | isCID Energy (eV) | 0.0 |
| **Spectrum Setting** | | Funnel 2 RF (Vpp) | 400.0 |
| Rate mode | Summation | Multipole RF (Vpp) | 400.0 |
| Rate value | 112 | **Collision cell** | |
| **Laser Settings** | | Collision Energy (eV) | 10.0 |
| Burst | 1 | Collision RF (Vpp) | 720.0 |
| Shots | 15 | **Quadrupole** | |
| Frequency (Hz) | 10000 | Ion energy (eV) | 5.0 |
| Laser power (%) | 80 | Low mass (*m/z*) | 50.0 |
| Application | Custom | **Focus pre-TOF** | |
| Power boost | 0.0 | Transfer time (µs) | 60.0 |
| Smart beam | On | Pre-pulse storage (µs) | 5.0 |
| Scan range (X & Y) (µm) | 16.0 | **Detection** | |
| Resulting field size (X &Y) (µm) | 5.0 | High sensitivity detection/ focus mode | Off |

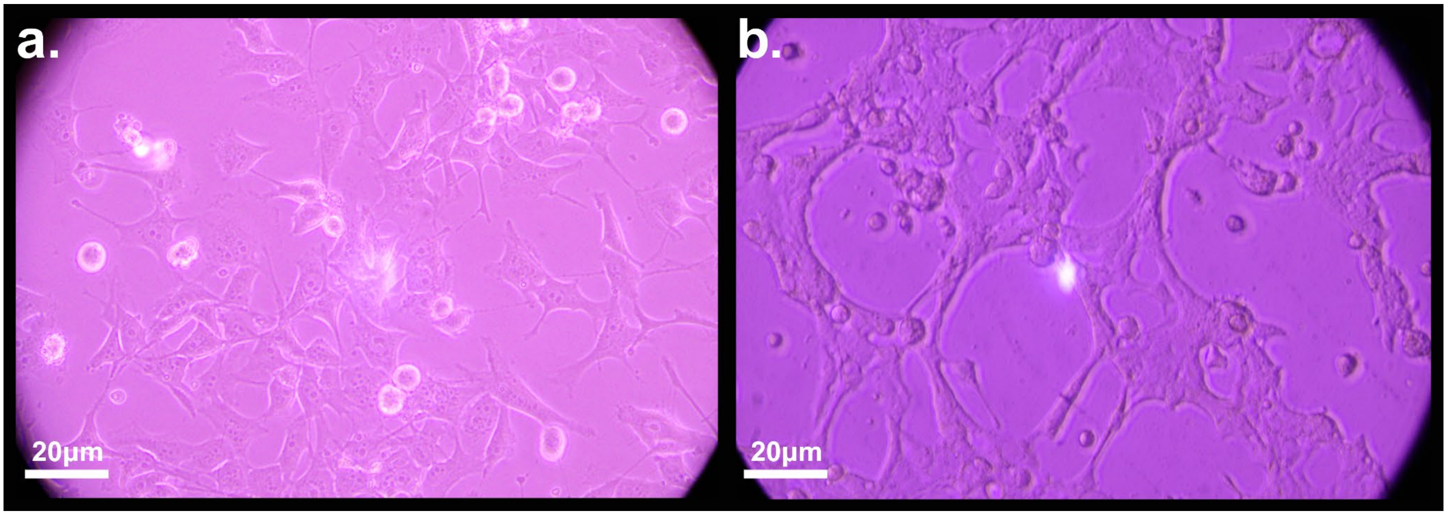

**Figure S2: OVCAR8-RFP cell morphology.** OVCAR8-RFP cells cultured in a T75 flask **(a)** were used as the morphological reference for comparison with OVCAR8-RFP cells adhered to the surface of the hydrogel **(b)**. Images were acquired using a Laxco SLi3Pro microscope equipped with a SeBa Cam Cool 1.7 MP camera (S/N: C2211211201, USB 3.0, DC 12V/3A) and a 45 mm LBD filter. Images were captured and color-corrected using SeBa View software.

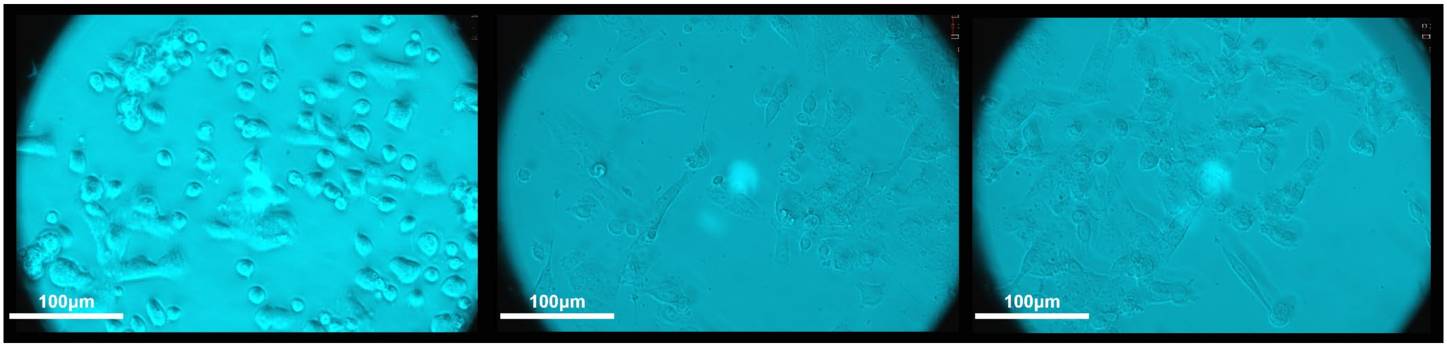

**Figure S3:** ***Evaluation of OVCAR8-RFP cell adhesion and proliferation on MaxGel™ECM functionalized hydrogels over 48 hours*.** Conditions composed from 100% MaxGel™ ECM were further monitored for an additional 24 hours beyond the initial incubation period. After the first 24 hours *(left)*, adherent cells were observed on the hydrogel surface. After 48 hours of total incubation (*middle and right*), a marked increase in cell adhesion and proliferation was observed, with cells displaying well-spread, polygonal morphologies indicative of healthy attachment. All samples were maintained at 37 °C in a humidified incubator with 5% $CO_2$. Images were acquired using a Laxco SLi3Pro microscope equipped with a SeBa Cam Cool 1.7 MP camera (S/N: C2211211201, USB 3.0, DC 12V/3A) and a 45 mm LBD filter. Images were captured and color-corrected using SeBa View software.

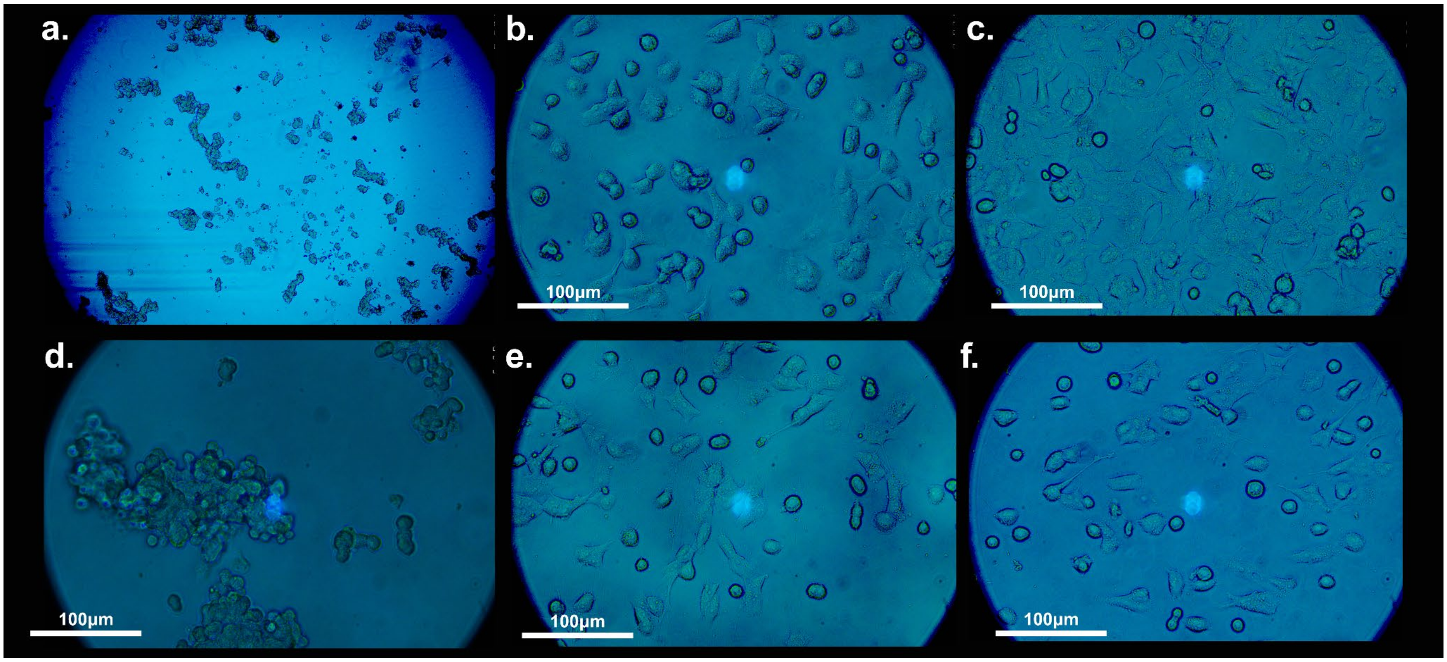

**Figure S4:** ***Poly-D-lysine rehydration and fibronectin functionalization of hydrogels.*** Hydrogels (30 mm diameter) were desiccated and rehydrated by submersion in 2 mL of 0.1 mg/mL PDL. Excess PDL was removed, and samples were air-dried for 10–15 minutes prior to sulfo-SANPAH activation and subsequent fibronectin coating as the cell-adhesion ligand. **(a)** Blank hydrogel control (4× objective). OVCAR8-RFP cells formed aggregates due to the absence of surface functionalization, indicating poor substrate adhesion and increased cell–cell interactions. **(d)** Hydrogels treated with sulfo-SANPAH only (no protein). Cells exhibited minimal surface attachment, high displacement after mechanical perturbation, and predominantly rounded morphologies, consistent with elevated cellular stress and poor adhesion. **(b,e)** Hydrogels functionalized with sulfo-SANPAH followed by 300 µg/mL fibronectin in 1× PBS. Moderate cell adhesion was observed. **(c,f)** Hydrogels functionalized with sulfo-SANPAH followed by 450 µg/mL fibronectin in 1× PBS. Robust cell adhesion with well-spread, polygonal morphologies was observed. Images were acquired using a Laxco SLi3Pro microscope equipped with a SeBa Cam Cool 1.7 MP camera (S/N: C2211211201, USB 3.0, DC 12V/3A) and a 45 mm LBD filter. Images were captured and color-corrected using SeBa View software.

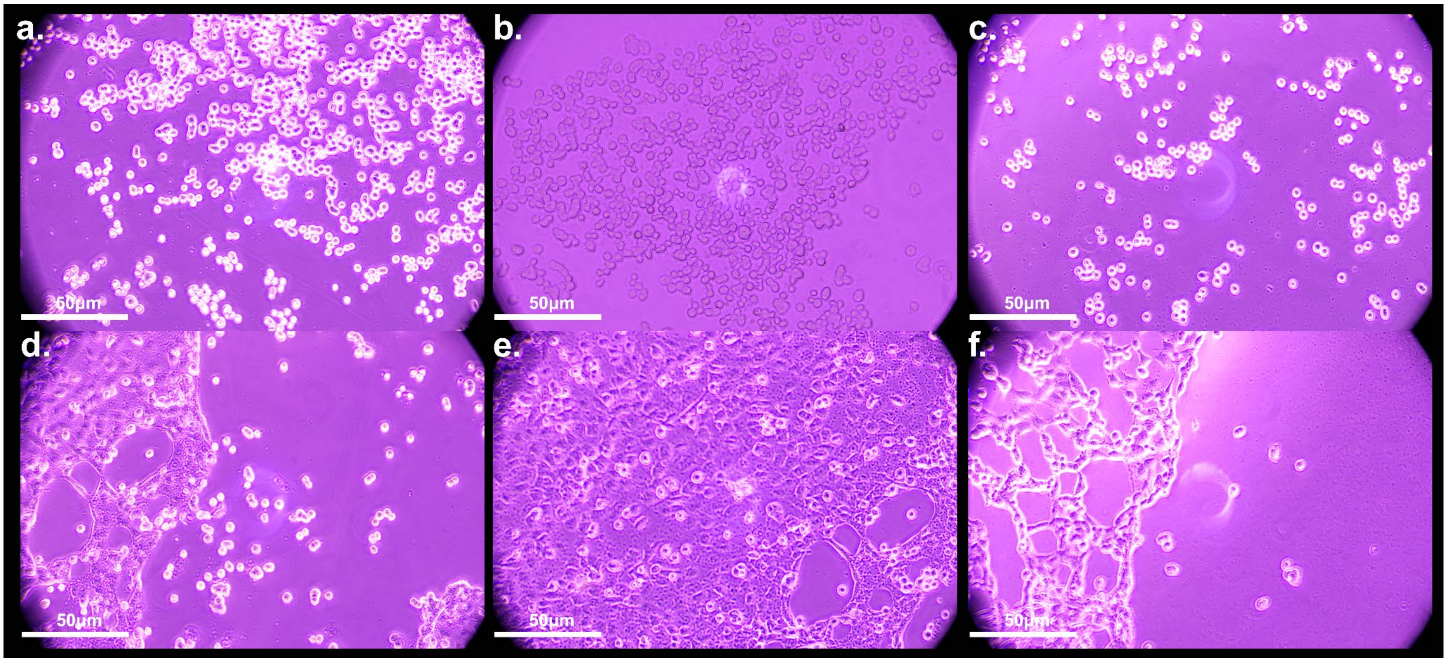

**Figure S5:** ***Effect of fibronectin incubation temperature on OVCAR8-RFP cell adhesion and morphology.*** To further enhance OVCAR8-RFP cell adhesion to the hydrogel substrate and ensure stable attachment during mechanically induced stress, fibronectin incubation temperature was evaluated. All hydrogels were prepared and functionalized under identical conditions using sulfo-SANPAH followed by fibronectin coating; PDL was not used in these experiments. The only variable was the temperature at which fibronectin was incubated on the hydrogel surface. In the top row **(a-c)**, hydrogels were incubated with fibronectin at 37 °C, whereas in the bottom row **(d-f)**, fibronectin incubation was performed at 4 °C. Each image represents an independent biological replicate(n=3). While OVCAR8 cells adhered under both conditions, clear differences in cellular morphology were observed. Hydrogels incubated at 37 °C **(a-c)** supported cell adhesion but predominantly exhibited a rounded or circular morphology. In contrast, hydrogels incubated at 4 °C **(d-f)** showed robust adhesion with polygonal cell morphology closely resembling that observed for OVCAR8 cells cultured on standard tissue-culture flasks (T75 flasks) forming networks within the gel. Images were acquired using a Laxco SLi3Pro microscope equipped with a SeBa Cam Cool 1.7 MP camera (S/N: C2211211201, USB 3.0, DC 12V/3A) and a 45 mm LBD filter. Images were captured and color-corrected using SeBa View software.

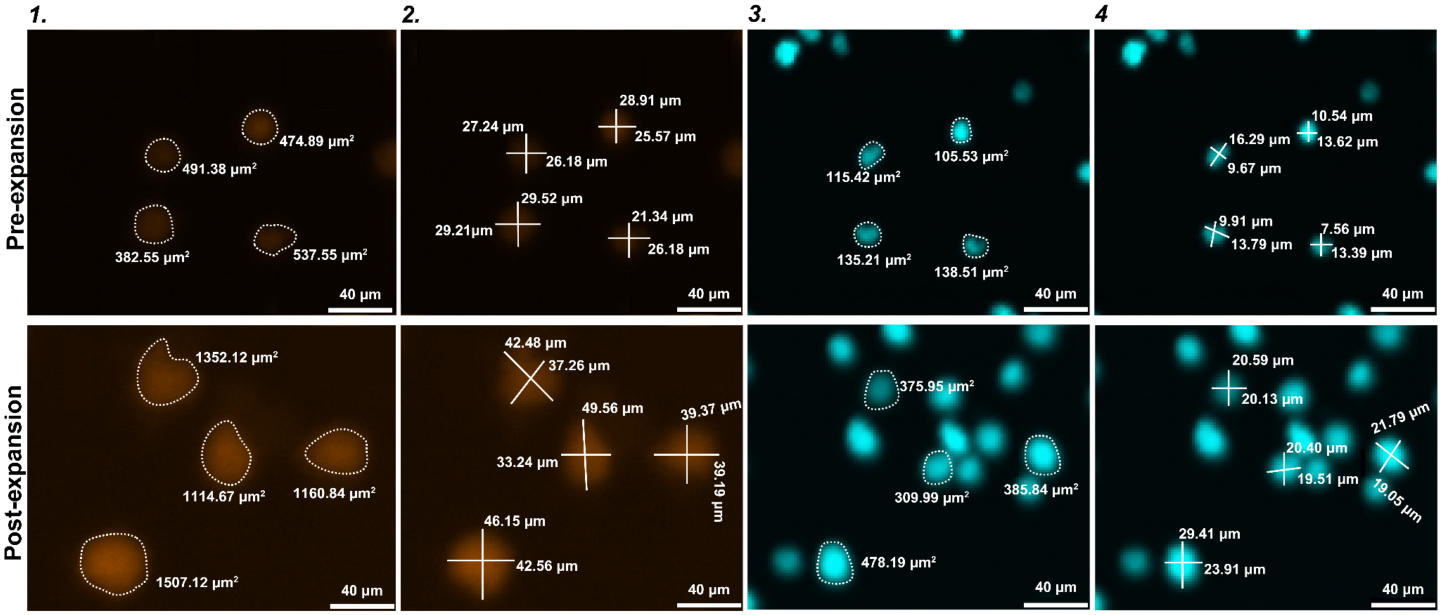

**Figure S6:** ***Evaluation of pre- and post-expanded OVCAR8-RFP cells using ImageJ (fiji).*** Additional analysis was performed on pre- and post-expanded OVCAR8-RFP cells to assess potential size changes induced by the expansion process. Both whole-cell and nuclear dimensions were evaluated using ImageJ (fiji). Cell and nucleus measurements were obtained by **(1,3)** quantifying area (µm²) and **(2,4)** diameter (µm) from fluorescence images. Measurements were collected from multiple regions of each sample to ensure representative sampling across conditions, enabling direct comparison between pre-expansion and post-expansion states. Cell area measurements were performed manually and reliably segment whole-cell areas. In contrast, nuclear area measurements were acquired automatically using the particle analysis function, as the nuclei could be consistently identified and segmented by the software.

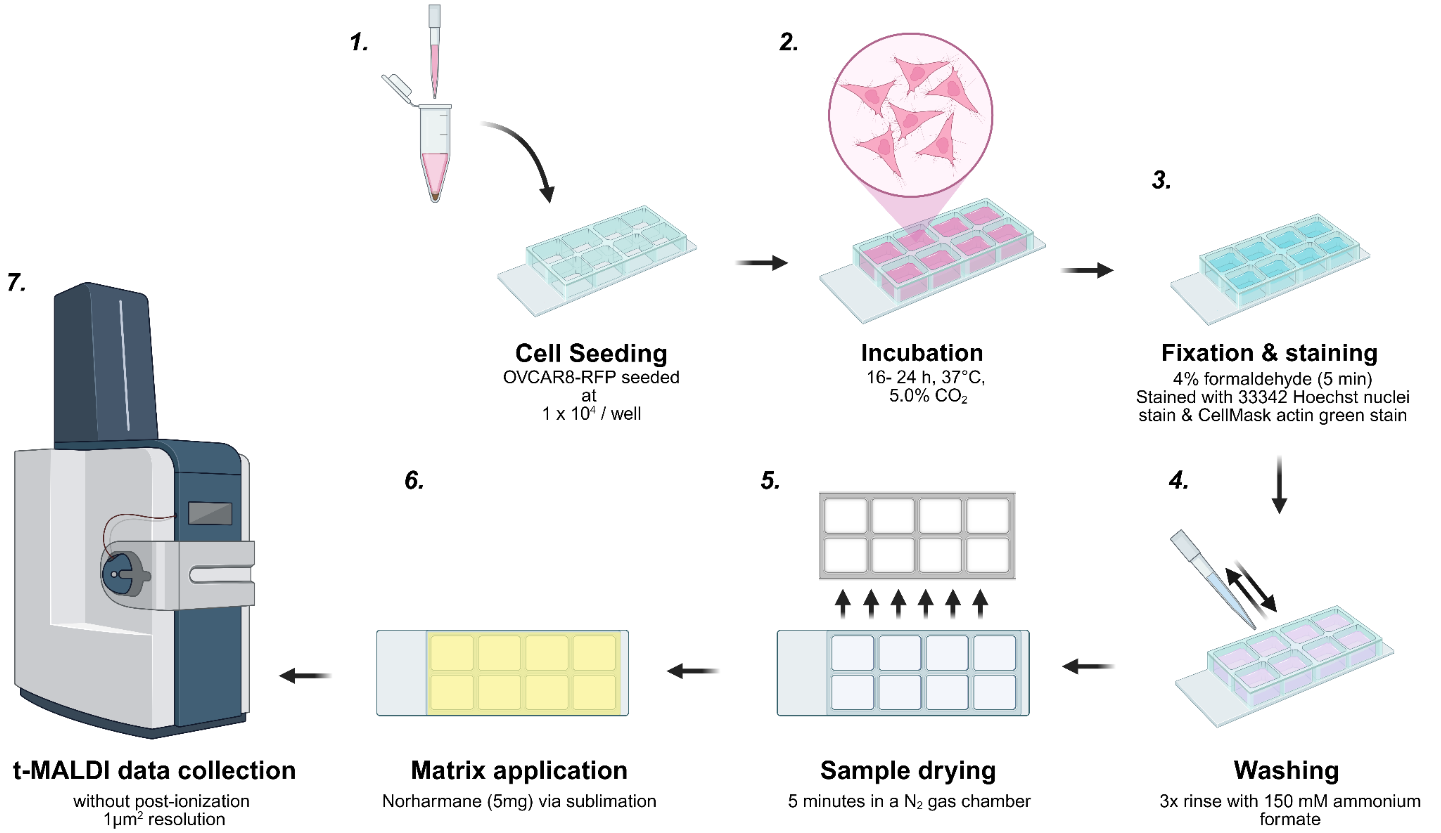

**Figure S7:** ***Sample preparation of transmission mode geometry-MALDI for single cell imaging*.** **(1)** In a sterile Millicell EZ slide add 500μL of DMEM 1x media, and add 10,000 of OVCAR8-RFP cells per well. **(2)** slides with cells are allowed to incubate over a 16-24 hour period allowing the cells to proliferate. **(3)** Media is removed from the wells, and rinsed with 500μL PBS 1x three times followed by fixation with 500 μL 4% formaldehyde. 250 μL 33342 Hoechst nuclei stains are added per well and left to rest from direct light for 5 minutes. Followed with a 500 μL PBS 1x rinse. Once the PBS 1x is removed, **(4)** rinse 3x with 150 mM of ammonium formate. Excess ammonium formate was removed to ensure low crystal formation during **(5)** during the drying process the 8 well chambers are removed and placed in as nitrogen gas chamber constant flow of 5 minutes. **(6)** followed by a matrix application of norharmane via sublimation to ensure small matrix crystals below the raster size and even energy distribution during sample collection. **(7)** data collection with an in-house built t-MALDI-MSI allowing for highly detailed resolution of single cells. Figure adapted from Bien et al.[1]

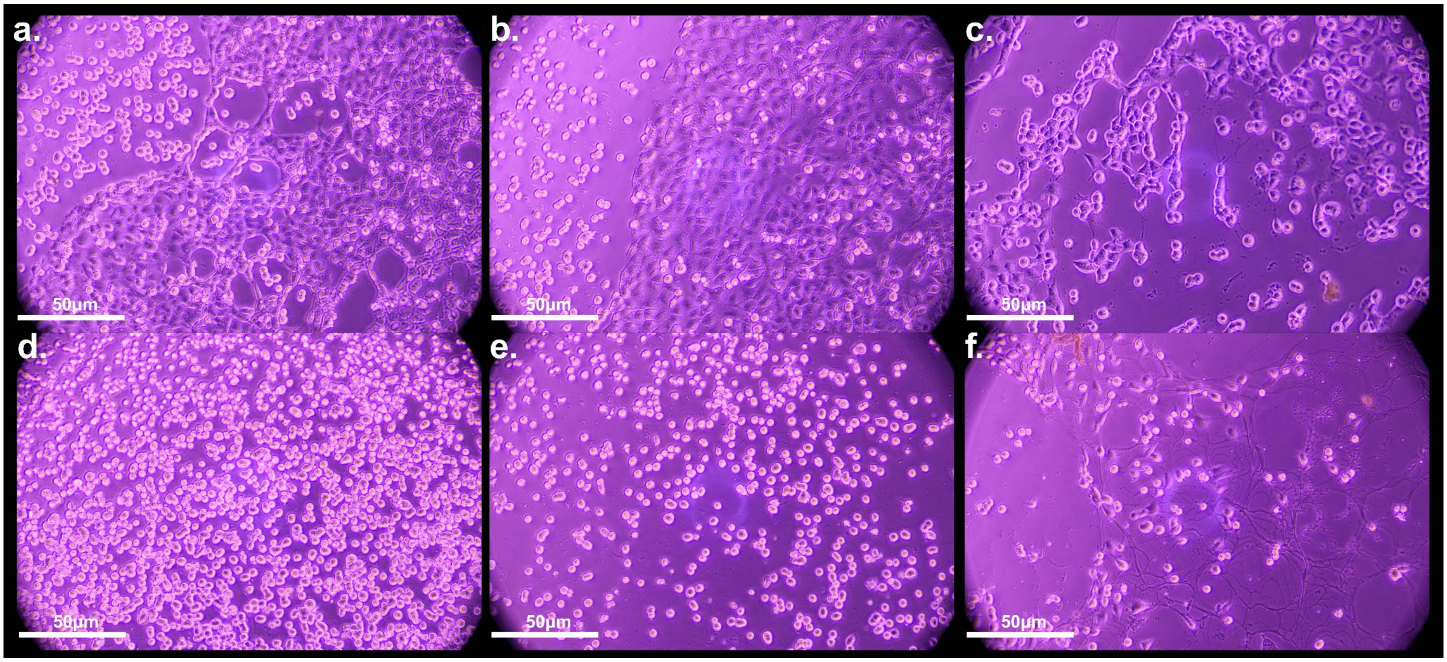

**Figure S8:** ***Serum-treated vs. serum-starved cell morphology.*** To probe metabolic stress and potential flux, OVCAR8-RFP cells were subjected to serum starvation (growth factor deprivation) to evaluate their viability and behavior on the hydrogel substrate. Panels **(a-c)** show cells cultured in the presence of fetal bovine serum, while panels **(d-f)** show serum-starved conditions. A clear morphological shift was observed, with serum-treated cells exhibiting a polygonal, adherent phenotype, whereas serum-starved cells adopted a more rounded morphology. This transition is consistent with a stress response, where cells reduce spreading and cytoskeletal activity to conserve energy under nutrient-deprived conditions. Images were acquired using a Laxco SLi3Pro microscope equipped with a SeBa Cam Cool 1.7 MP camera (S/N: C2211211201, USB 3.0, DC 12V/3A) and a 45 mm LBD filter. Images were captured and color-corrected using SeBa View software.

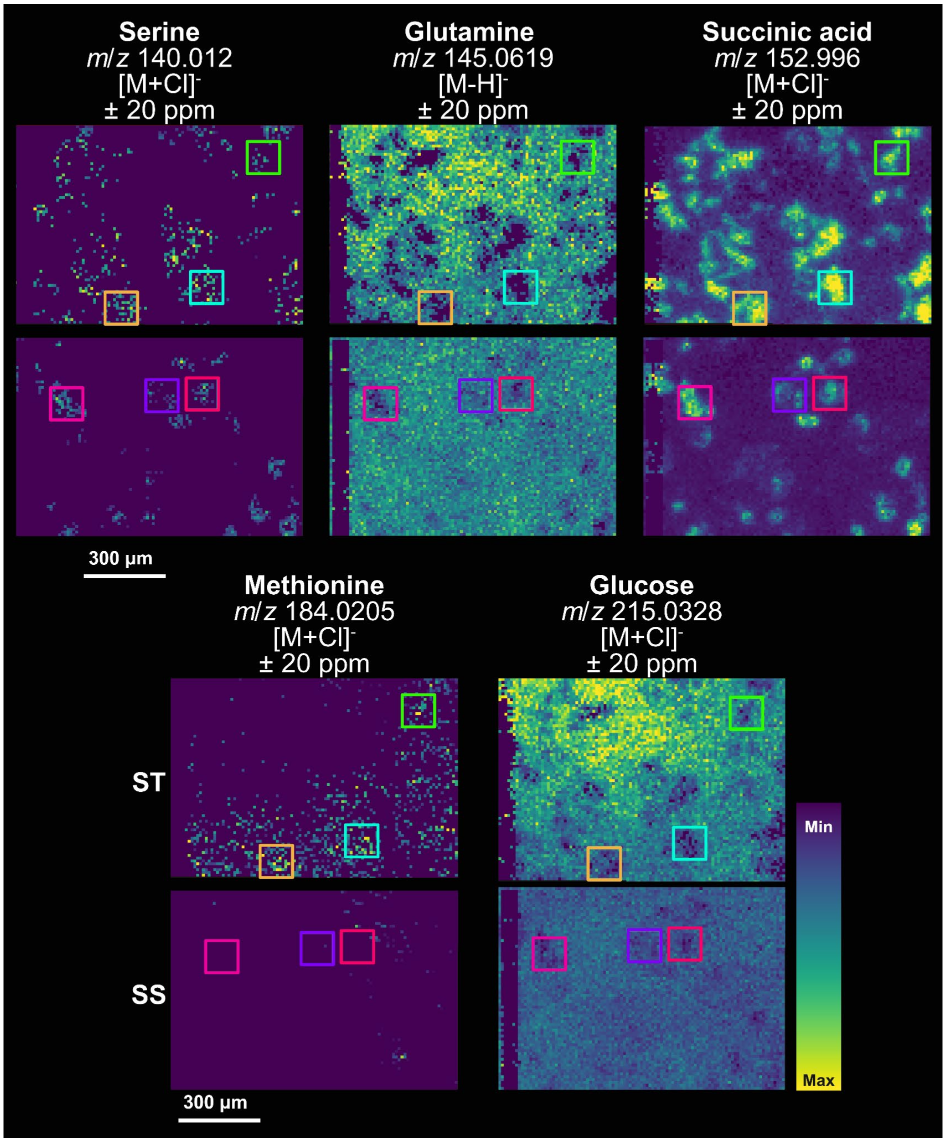

**Figure S9:** ***Evaluation of OVCAR8-RFP metabolic flux cells under serum treated (ST) and serum starved conditions within regions of interest (ROI).*** Marked in different colors (orange, cyan, green, pink, purple, and magenta) are ROI's that were utilized for log fold 2 change (Fig. S14).

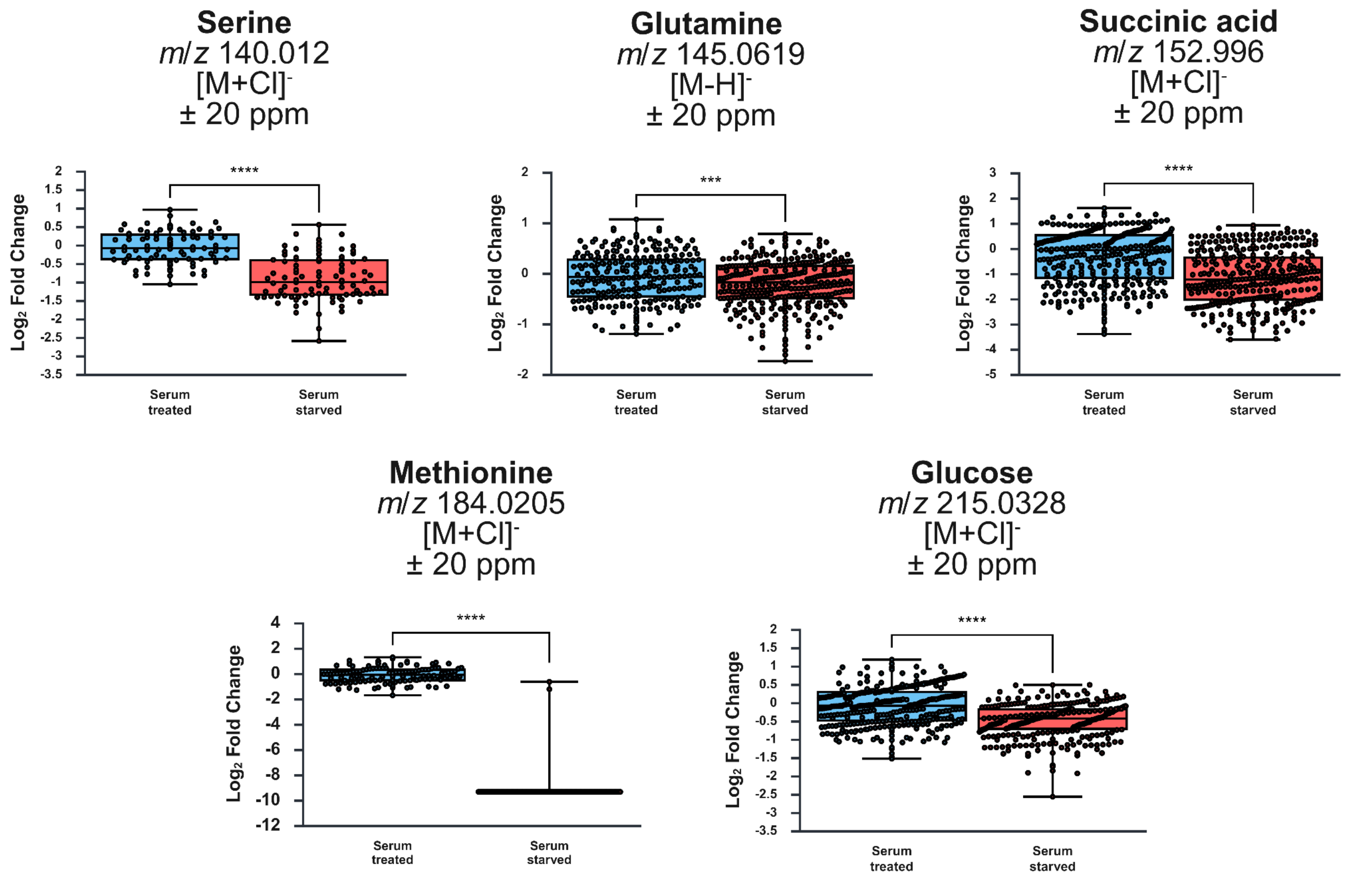

**Figure S10:** ***Evaluation of metabolic flux serum-treated vs serum-starved OVCAR8-RFP.*** MALDI-MSI data were processed using SCiLS Lab, and ion images were normalized to the total ion current (TIC). Prior to the TExMS workflow, serum-starved cells exhibited a more spherical morphology compared to the polygonal morphology observed in serum-treated controls. This morphological shift likely reflects the removal of FBS-derived growth factors, proteins, and nutrients necessary for proliferation and adhesion, causing cells to transition from a proliferative state toward a survival or stress-response state. Correspondingly, distinct differences in metabolic profiles were observed between the two conditions.

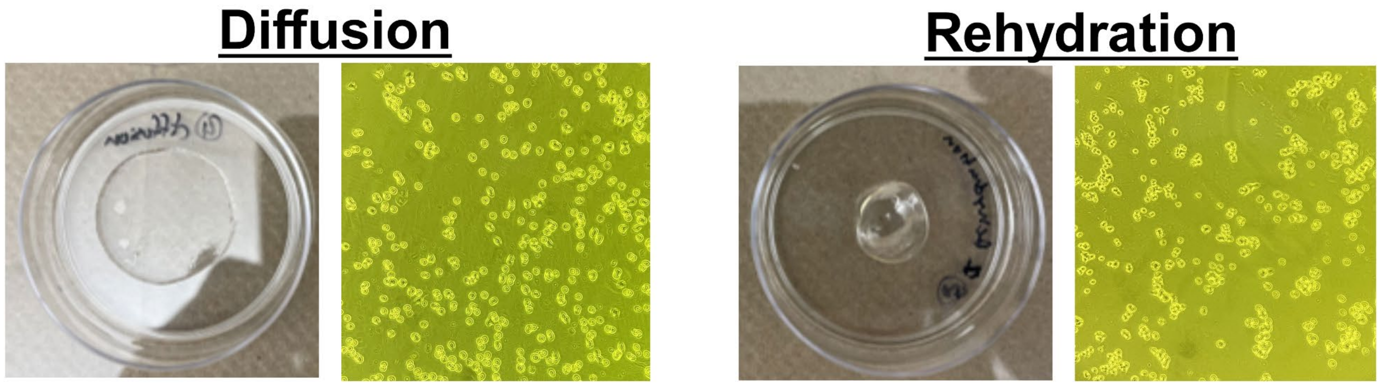

**Figure S11: Comparison of surface functionalization strategies for promoting cell adhesion on hydrogels.** Two 30 mm diameter hydrogels were prepared using different approaches to promote cell adhesion. Hydrogel 1 (*left*) was coated with 1 mL of 0.1 mg/mL PDL and incubated at room temperature for 10 minutes. Excess PDL was then removed, and the hydrogel was rinsed with sterile MilliQ $H_2O$. For Hydrogel 2 (*right*), the sample was first desiccated in an oven at 35 °C for 2 hours to remove excess water from the hydrogel network. Once the hydrogel appeared dehydrated, 5 mL of 0.1 mg/mL PDL was added and allowed to incubate for 10 minutes to rehydrate the network with PDL, followed by rinsing with sterile MilliQ $H_2O$. Both functionalization strategies resulted in comparable levels of cell adhesion to the hydrogel surface. However, cells exhibited a predominantly spherical morphology under both conditions, in contrast to the flattened, polygonal morphology typically observed on rigid substrates. Images were collected on a IPhone 12 pro max camera.

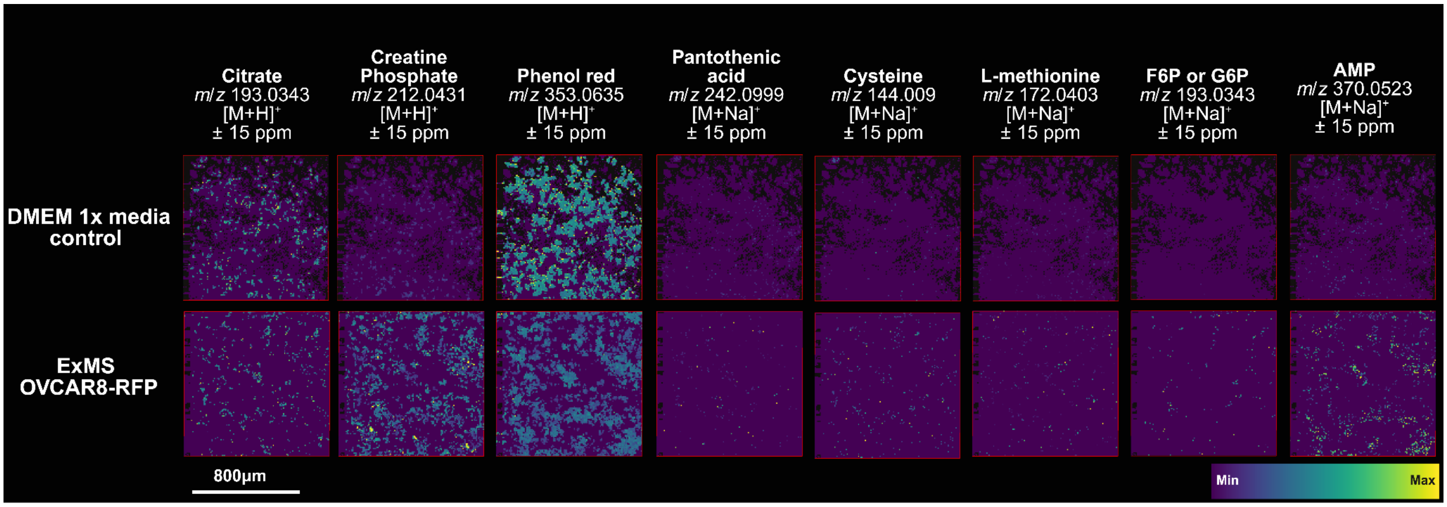

**Figure S12:** ***TExMS 03 matrix optimization.*** The CHCA matrix concentration was reduced to 5 mg/mL, corresponding to a final matrix density of 0.0036 $mg/mm^2$ and a laser power of 70%. In addition, the matrix application temperature was increased from 70 °C to 90 °C, promoting the formation of smaller and more homogeneous matrix crystals. These adjustments reduced hotspot formation associated with matrix oversaturation and improved signal uniformity. Under these optimized conditions, small molecules were successfully putatively annotated in positive ion mode.

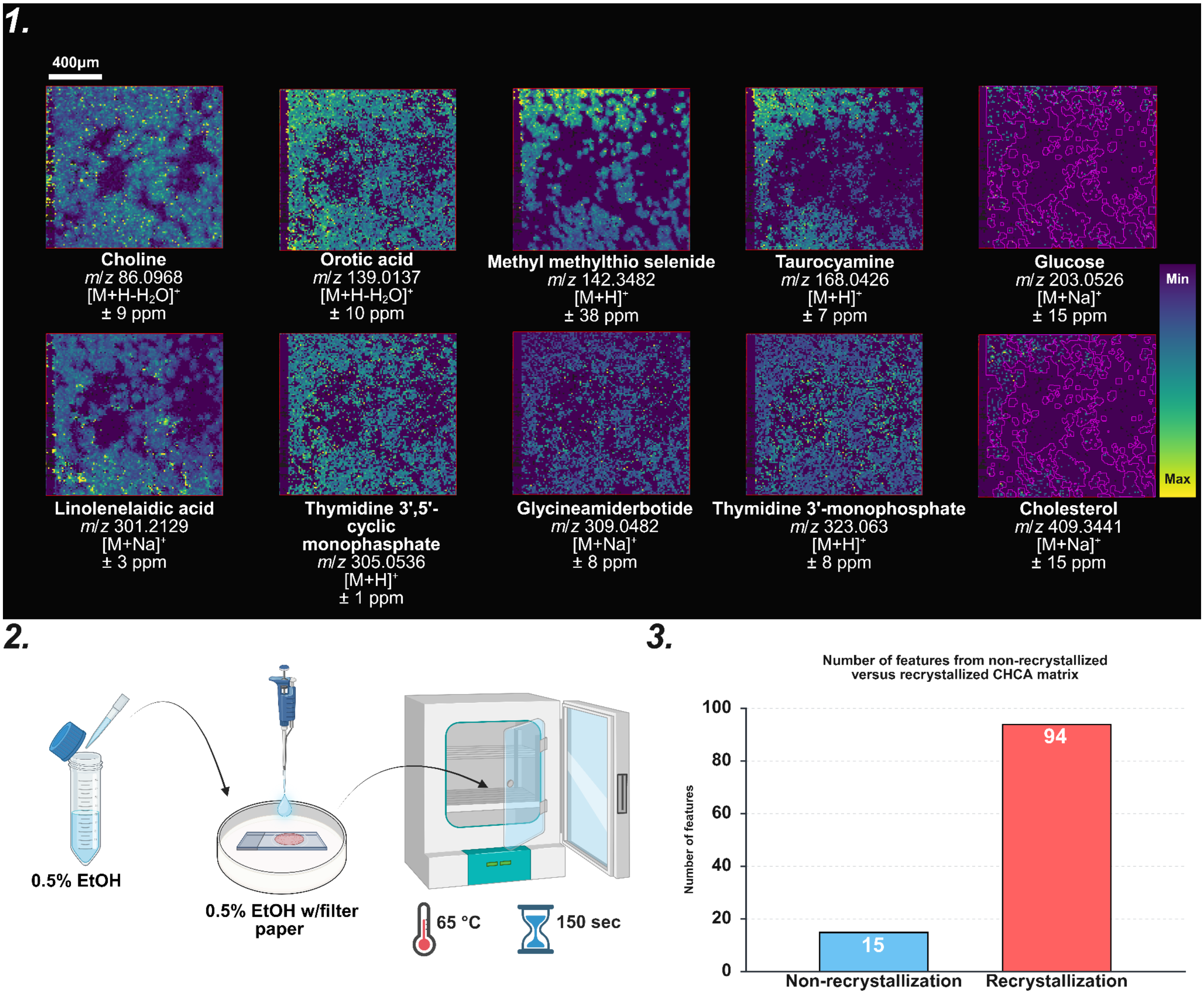

**Figure S13:** ***TExMS 05 matrix recrystallization and topography optimization.*** To improve sample flatness and maintain optimal topography, an 18 mm circular coverslip was incorporated beneath the sample **(1)**. To further enhance ion signal intensity and spatial resolution, CHCA matrix density: 0.0051 mg/mm²) recrystallization was performed. Under optimized conditions (laser power: 65%), an increased number of detectable features was observed. **(2)** For recrystallization, a 0.5% ethanol (EtOH) solution was used to generate a vapor environment. A 90 mm Petri dish containing an 85 mm filter paper was prepared, and 500 µL of the EtOH solution was applied to the filter paper without disturbing the sample. The chamber was then heated at 65°C for 150 seconds. **(3)** This approach resulted in an increased number of spectral features compared to non-recrystallized samples; however, of the 94 detected features, only 18 were successfully annotated. Additionally, Hoechst 33342 nuclear stain was introduced during this optimization to provide a distinct m/z marker for cell localization. However, no detectable ion signal corresponding to the stain was observed under these experimental conditions.

**Table S4:** Additional putative annotations of TExMS 05 recrystallized data.

| Observed MSI signal | Calculated exact mass | Putative annotation | Adduct | ± ppm error |
|---|---|---|---|---|
| 167.0081 | 167.0068 | 5,6-Dihydroxyuracil | $[M+Na]^+$ | 11 |
| 168.0426 | 168.0442 | Taurocyamine | $[M+H]^+$ | 7 |
| 201.0618 | 201.0639 | Endostatin | $[M+Na]^+$ | 8 |
| 242.0169 | 242.0451 | 3-Mercaptolactate-cysteine disulfide | $[M+H]^+$ | 7 |
| 273.1804 | 273.1814 | N-octanoylglutamine | $[M+H]^+$ | 2 |
| 299.1955 | 299.1987 | 19-Norandrosterone | $[M+Na]^+$ | 9 |
| 301.2129 | 301.2143 | Linolenelaidic acid | $[M+Na]^+$ | 3 |
| 306.0527 | 306.0491 | Cytidine 2',3'-cyclic phosphate | $[M+H]^+$ | 14 |
| 427.3487 | 427.3551 | (22alpha)-hydroxy-cholestanol | $[M+Na]^+$ | 15 |

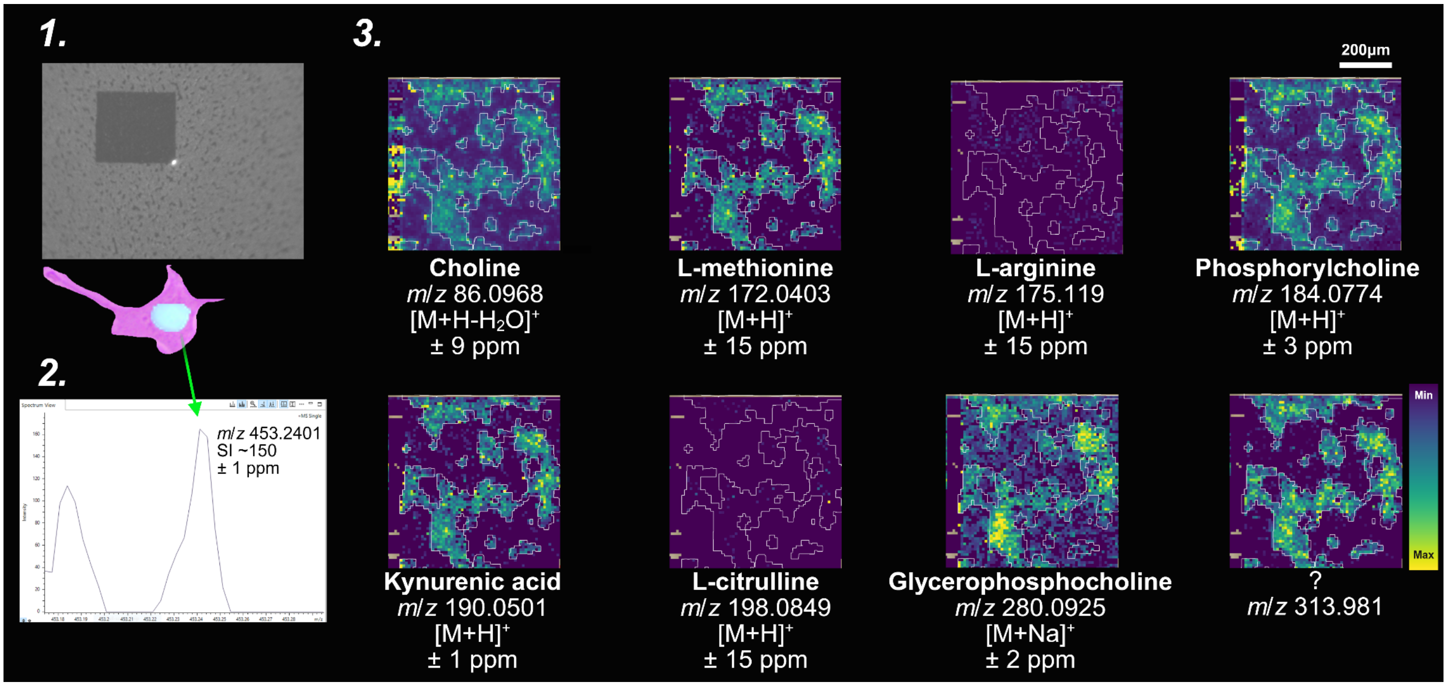

**Figure S14:** ***TExMS 06 optimization for ionization efficiency and cell localization.*** The incorporation of an 18 mm circular coverslip **(1)** improved sample flatness, resulting in a more uniform topography. Under these conditions (laser power: 70%), CHCA matrix matrix density: 0.0018 mg/mm$^2$) deposition produced smaller, more homogeneous crystals. In a subsequent attempt **(2)**, a signal at m/z 453.24—attributed to the Hoechst 33342 nuclear stain—was detected; however, it remained at noise-level intensity and was not reliably observed during MSI acquisition. Despite the lack of a robust Hoechst signal, overall sample ionization **(3)** was consistent and reproducible.

**Table S5:** HTX TM Sprayer parameters with TExMS 03-06 (CHCA) experiments.

| | | | |
|---|---|---|---|
| **Matrix** | CHCA | **Velocity (mm/min)** | 1100 |
| **Concentration (mg/mL)** | 5 | **Pressure (psi)** | 10 |
| **Solvent (%)** | ACN:$H_2O$ (9:1) 0.1% TFA | **Track Spacing** | CC |
| **Temp(°C)** | 90 | **Gas flow rate (L/min)** | 3 |
| **Passes** | 12 | **Drying time (s)** | 0 |
| **Flow rate (mL/min)** | 0.2 | **Nozzle Height (mm)** | 40 |

**Table S6:** MS Ion-optics parameters for TExMS (CHCA) experiments.

| **MS Setting** | | **Tune** | |
|---|---|---|---|
| Scan begins (m/z) | 50 | MALDI Plate offset (V) | 75.0 |
| Scan ends (m/z) | 1200 | Deflection 1 delta (V) | 75.0 |
| Ion polarity | Positive | Funnel 1 RF (Vpp) | 150.0 |
| Scan mode | MS | isCID Energy (eV) | 0.0 |
| **Spectrum Setting** | | Funnel 2 RF (Vpp) | 200.0 |
| Rate mode | Summation | Multipole RF (Vpp) | 200.0 |
| Rate value | N/A | **Collision cell** | |
| **Laser Settings** | | Collision Energy (eV) | 10.0 |
| Burst | 1 | Collision RF (Vpp) | 350.0 |
| Shots | 1000 | **Quadrapole** | |
| Frequency (Hz) | 5000 | Ion energy (eV) | 5.0 |
| Laser power (%) | 65-70 | Low mass (*m/z*) | 50.0 |
| Application | Custom | **Focus pre-TOF** | |
| Power boost | 0.0 | Transfer time (µs) | 55.0 |
| Smart beam | On | Pre-pulse storage (µs) | 5.0 |
| Scan range (X & Y) (µm) | 6.0 | **Detection** | |
| Resulting field size (X &Y) (µm) | 10.0 | High sensitivity detection/ focus mode | On |

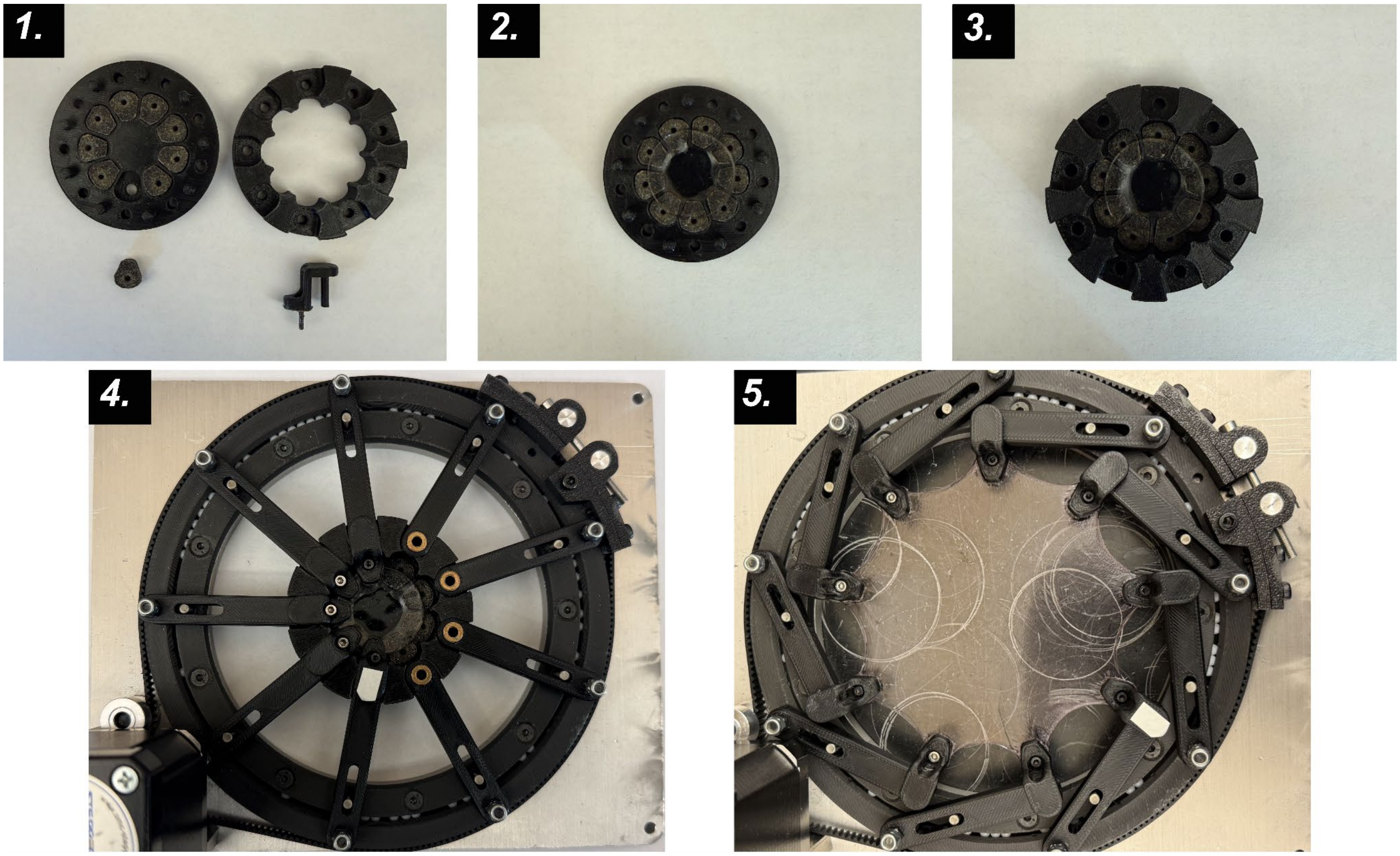

**Figure S15:** ***Setting up samples for tensile stretching.*** The iris stretcher requires a custom 3D-printed mounting apparatus to securely grip and mechanically stretch the hydrogel. **(1)** The mounting apparatus consists of upper and lower support components, while each gripper is composed of three parts: a lower gripper, an upper gripper, and an M3 screw. **(2)** A 30 mm diameter hydrogel is centered on the mounting apparatus, avoiding the screw holes. Proper positioning prevents the hydrogel from becoming entangled with the screws while providing sufficient contact area to minimize localized stress and reduce the risk of tearing during expansion. **(3)** Once the hydrogel is centered, the upper support component is positioned over the hydrogel to secure the iris stretcher arms. **(4)** The upper gripper components are then attached and fastened with M3 screws. **(5)** After the assembly is fully secured, the hydrogel is mechanically expanded to the desired expansion factor and stretching speed.
